# Mapping the topographic organization of the human zona incerta using diffusion MRI

**DOI:** 10.1101/2024.09.05.610266

**Authors:** Roy AM Haast, Jason Kai, Alaa Taha, Violet Liu, Loxlan W Kasa, Greydon Gilmore, Maxime Guye, Ali R. Khan, Jonathan C. Lau

## Abstract

The zona incerta (ZI) is a deep brain region originally described by Auguste Forel as an “immensely confusing area about which nothing can be said.” Despite the elusive nature of this structure, mounting evidence supports the role of the ZI and surrounding regions across a diverse range of brain functions and as a candidate target for neuromodulatory therapies. Using *in vivo* diffusion MRI and data-driven connectivity, we identify a topographic organization between the ZI and neocortex. Specifically, our methods identify a rostral-caudal gradient predominantly connecting the frontopolar and ventral prefrontal cortices with the rostral ZI, and the primary sensorimotor cortices with the caudal ZI. Moreover, we demonstrate how clustering and gradient approaches build complementary evidence including facilitating the mapping of a central region of the ZI, connected with the dorsal prefrontal cortex. These results were shown to be replicable across multiple datasets and at the individual subject level, building evidence for the important role of the ZI in mediating frontal lobe-associated tasks, ranging from motor to cognitive to emotional control. Finally, we consider the impact of this topographic organization on the refinement of neuromodulatory targets. These results pave the way for an increasingly detailed understanding of ZI substructures, and considerations for *in vivo* targeting of the ZI for neuromodulation.

## Introduction

Since its first description by Auguste Forel in 1877, the zona incerta (ZI) has remained one of the most elusive regions of the human brain. Located in the subthalamic region, the ZI is situated between the ventral thalamus, subthalamic nucleus (STN), and the red nucleus. This small gray matter region is surrounded by the H1 and H2 fields of Forel, the H field, as well as a number of important white matter pathways including the corticospinal, cerebellothalamic, and medial lemniscal tracts (Lau et al., 2020; Morel et al., 1997). Recent studies demonstrate that the ZI exhibits broad connections with the cortex, subcortical structures, brain stem, and spinal cord, resulting in theories regarding the role of the ZI as an integrative hub across a broad range of functions (Arena et al., 2024; Haber et al., 2023; Mitrofanis, 2005; Ossowska, 2020), including fear processing (Lin et al., 2023; Venkataraman et al., 2021; Zhou et al., 2021, 2018), sleep (Liu et al., 2017, 2011), attention (Chometton et al., 2021; Mitrofanis et al., 2004), and locomotion regulation (Garau et al., 2023; Kim et al., 2023; Liu et al., 2011; Sharma et al., 2018).

Based on observations in animal models, the ZI has been identified to consist of a unique molecular environment that is rich in terms of cellular and functional diversity (Chometton et al., 2017; Moriya and Kuwaki, 2020; Wu et al., 2023; Yang et al., 2022). The ZI has been determined to be predominantly composed of inhibitory GABAergic neurons with additional populations such as glutamatergic, dopaminergic, and melanin-concentrating hormone neurons, and upwards of 20 different neuron populations (Fratzl and Hofer, 2022; Mitrofanis et al., 2004; Watson et al., 2014). Based on these cytoarchitectural profiles, the ZI can be divided into 6 internal neuronal subregions in rodents: pars rostropolaris, or rostral ZI (rZI), pars dorsalis, or dorsal (dZI), pars ventralis, or ventral (vZI), pars magnocellularis, pars retropolaris, and pars caudalis (Kawana and Watanabe, 1981; Romanowski et al., 1985). The latter 3 sectors are later collectively known as the caudal ZI (cZI) (Arena et al., 2024; Nicolelis et al., 1995), whereas the dZI and vZI are commonly referred to as the central region in contemporary literature (Monosov et al., 2022; Watson et al., 2014). Furthermore, these ZI subregions exhibit differential connectivity patterns with the cortex, supporting a critical role of cortico-incertal connections for brain-wide communication across a range of functions (Barthó et al., 2002; Lin et al., 2023; Venkataraman et al., 2021; Wang et al., 2024; L. Zhang et al., 2022; Zhang and van den Pol, 2017; Zhou et al., 2021). The extent to which rodents recapitulate findings in the human ZI remains poorly understood due to substantial phylogenetic differences. Not only is the human cortex much larger in size, but it also has regions with no corresponding homologues in mice. Genetic differences, such as primate-specific neural progenitors driving cortical development, and distinct functional connectivities further highlight cross-species disparities. Therefore, direct research using primate data is essential to fully understand the ZI in humans.

Poor direct visualization of the ZI has contributed to the challenges with performing complementary analyses in humans. The cZI remains the most well-studied subregion in humans given its efficacy as a deep brain stimulation (DBS) target for Essential Tremor (ET) (Awad et al., 2020; Plaha et al., 2011; Shapson-Coe et al., 2021) and Parkinson’s Disease (PD) (Bingham and McIntyre, 2024; Blomstedt et al., 2018; Mostofi et al., 2019; Plaha et al., 2006). However, the lack of visibility using standard magnetic resonance imaging (MRI) sequences has led the surgical target to be more conventionally referred to as the “posterior subthalamic area” given it has been unclear what structure is being targeted (Blomstedt et al., 2010). Manual segmentations in stereotactic (MNI) space provide an avenue for estimating the ZI location and, when combined with diffusion MRI (dMRI), point to growing evidence of functional duality with the rZI putatively modulating obsession-compulsive behavior and the cZI modulating tremor (Saluja et al., 2024). Studies in non-human primates have provided some of the critical evidence validating these findings in the rostral and caudal ZI using dMRI with validation via tract tracing methods (Haber et al., 2023). These studies build evidence for dMRI as a viable method for characterizing cortical-incertal connections *in vivo*.

Building on the aforementioned work, we aim to provide a refined analysis of the topographic organization of the human ZI by mapping its cortical, i.e., *cortico-incertal*, connections. Our approach includes both “classical” discrete clustering via spectral clustering and the use of diffusion map embedding to identify continuous connectivity gradients. Diffusion map embedding, a method for mapping transitions in structural connectivity, provides nuanced views of connectivity differences that might resemble those observed in animal models better (Margulies et al., 2016; Royer et al., 2024). We combine both computational neuroanatomy techniques with our previous *in vivo* labeling of the human ZI, based on ultra-high field (7 Tesla; 7T) MRI T_1_ mapping (Lau et al., 2020), and high-quality dMRI datasets from the Human Connectome Project (HCP) (Van Essen et al., 2012). Our structural connectivity workflow replicates established protocols that have previously been used to study other subcortical regions (Lambert et al., 2012). We demonstrate that our ZI structural connectivity maps (1) recapitulate known subregions from previous rodent and primate studies, (2) are replicable across field strengths and individual subjects, (3) can be linked to previously established cortical hierarchies and functional specializations, and (4) offer a proof-of-concept for personalizing neuromodulation strategies.

## Results

### Characterizing human cortico-incertal connectivity using *in vivo* tractography

We reconstructed cortico-incertal structural connectivity using *in vivo* probabilistic tractography based on high-resolution 7T dMRI HCP data (N=169) (Van Essen et al., 2012) and our previously validated ZI segmentation (Figure 1a-b) (Lau et al., 2020). The connections between the cortex and ZI were predominantly between the ZI and frontal lobe, representing approximately 65-75%, and secondarily connections were identified with the parietal lobe at 15-25%, with contributions identified to the temporal and occipital lobe were much more sparse (<5%) (Supplementary Figure 1). The majority of connections between the ZI and frontal lobe passed by way of the internal capsule.

**Figure 1.**
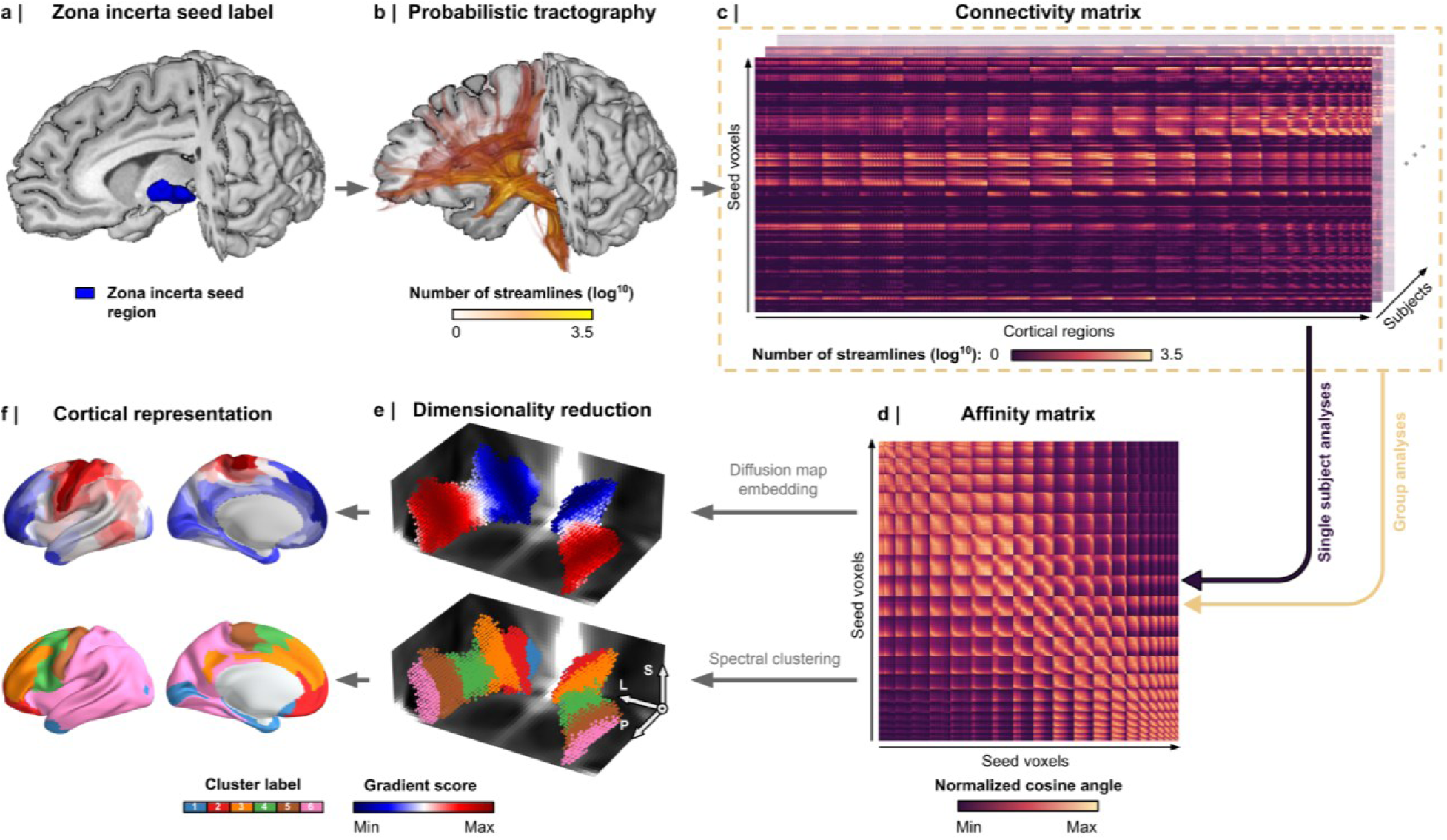
Cortico-incertal connectivity analysis workflow. (a) The zona incerta (ZI) seed region used for connectivity analyses. Diffusion MRI data were processed to reconstruct (b) streamlines via diffusion tractography and (c) connectivity matrices quantifying the number of streamlines between each ZI voxel and cortical region as defined by the HCP-MMP1.0 atlas. (d-f) ZI gradients and clusters were computed to illustrate the principal organizations of connectivity variability among ZI voxels. (d) Significant patterns were highlighted based on inter-voxel similarity using the normalized cosine angle. (e) Diffusion map embedding and spectral clustering were used to construct the gradients and clusters, respectively. (f) Gradient-weighted cortical maps were created by multiplying each row of the initial connectivity matrices with the corresponding principal gradient value, then averaging these rows to produce a single cortical representation of each gradient. A winner-takes-all approach was used to create a cortical map with areas color-coded according to their connectivity with the ZI clusters.

To investigate whether a more granular topography of cortico-incertal organization was present, the initial analysis of the group-average tractogram was refined by applying diffusion map embedding and spectral clustering to the group-average structural connectivity matrix. For each subject, cortico-incertal connectivity was represented as *M*-by-*N* matrices, where *M* corresponds to the voxels in the ZI (N=1981 for the left hemisphere and N=1901 for the right hemisphere) and *N* corresponds to the cortical parcels from the HCP multimodal parcellation (HCP-MMP1.0, N=180 per hemisphere). These matrices detail the streamline counts between the ZI seed voxels and cortical brain regions (Figure 1c). Diffusion map embedding and spectral clustering of the connectivity matrix, averaged across all subjects, subsequently yielded continuous gradients of connectivity differences among ZI voxels (ranked based on their explained variance) and discrete clusters of ZI voxels with similar structural connectivity patterns, respectively (Figure 1d-f).

### Mapping continuous gradients of cortico-incertal connectivity reveals a predominant rostral-caudal organization of the ZI

The diffusion map embedding results demonstrated a predominant differentiation along the rostral-caudal axis of the ZI (Figures 2a and 2b). The primary gradient (G1) extracted using diffusion map embedding distinguished between the rZI (blue) and cZI (red), while the secondary gradient (G2) identified the interface between them (red) and the rest (blue). G1 reflected a connectivity gradient from voxels highly connected to primary sensorimotor (high G1) to more anterior prefrontal cortical areas including the frontal pole (low G2, Figure 2c). G2 captured an additional gradient of ZI voxels connected to premotor and dorsolateral prefrontal areas (low G2) versus the rest of the brain (high G2). Note that additional gradients were computed (see Supplementary Figure 2a).

**Figure 2.**
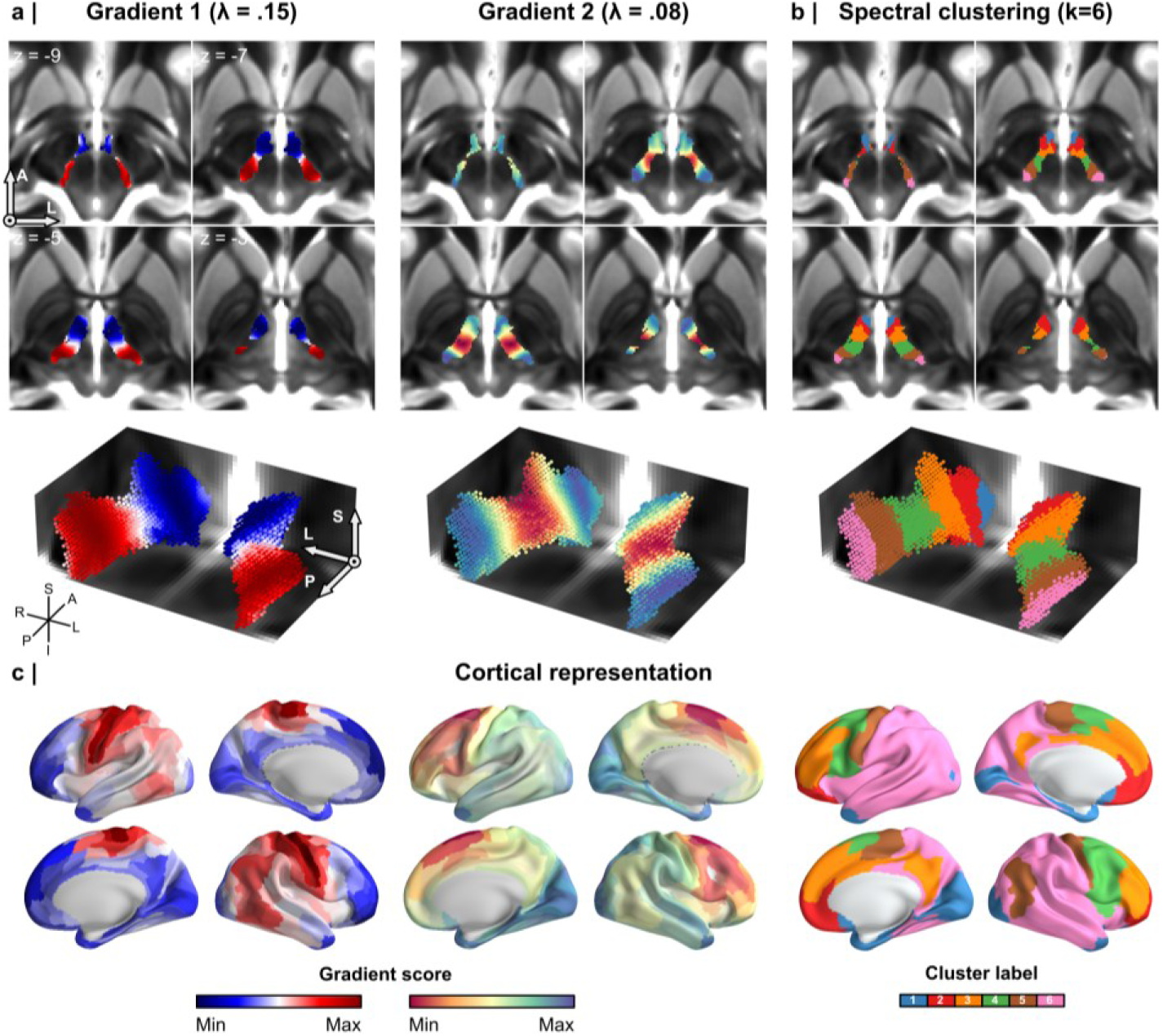
Cortico-incertal structural connectivity patterns. (a) The first two gradients of the zona incerta (ZI) based on structural connectivity shown using axial and 3D radiological views both revealed a rostral-caudal axis. (b) Similarly, spectral clustering shows a topographic organization of discrete clusters along a rostral-caudal axis, with cluster 1 positioned most rostrally. (c) Gradient-weighted cortical maps corresponding to gradients 1 and 2 and the spectral clustering results (left to right).

### Identifying discrete ZI subregions with differential cortico-incertal connectivity patterns

In parallel with the gradient approach, we selected a spectral clustering solution of k=6, that integrates cytoarchitectonic knowledge of the number of ZI sectors identified in rodent models (i.e., rZI, dZI, vZI and cZI) (Romanowski et al., 1985) with the added benefit of providing a more granular representation of the ZI given use cases include stereotactic targeting (see final section). Other cluster solutions were also computed (k=2-8, Supplementary Figure 2b). A composite visualization of the cluster-wise tractograms, illustrating the rostral-caudal organization of cortico-incertal connections, is displayed in Supplementary Figure 3, and their top 5% of streamlines are summarized in Supplementary Table 1.

Spectral clusters 1 to 6 followed the change in G1 values, where Cluster 1 represented the lowest G1 values and Cluster 6 the highest (Figure 2b). Looking at the cluster-wise tractograms (Figure 3), the rZI involving Clusters 1-2 were demonstrated to be structurally connected with the anterior and ventral prefrontal cortex including the frontal pole (BA10), orbitofrontal cortex (BA47), as well as the pregenual and subgenual components of the anterior cingulate cortex (ACC), consistent with studies using tract tracing in macaques and dMRI in humans (Haber et al., 2023; Saluja et al., 2024). The cZI (Clusters 5-6) was demonstrated to be highly connected to the primary motor cortex (BA04; shown in green in Supplementary Figure 1b), consistent with known literature in rodent models (Yang et al., 2022) as well as human and non-human primates (Haber et al., 2023; Saluja et al., 2024).

**Figure 3.**
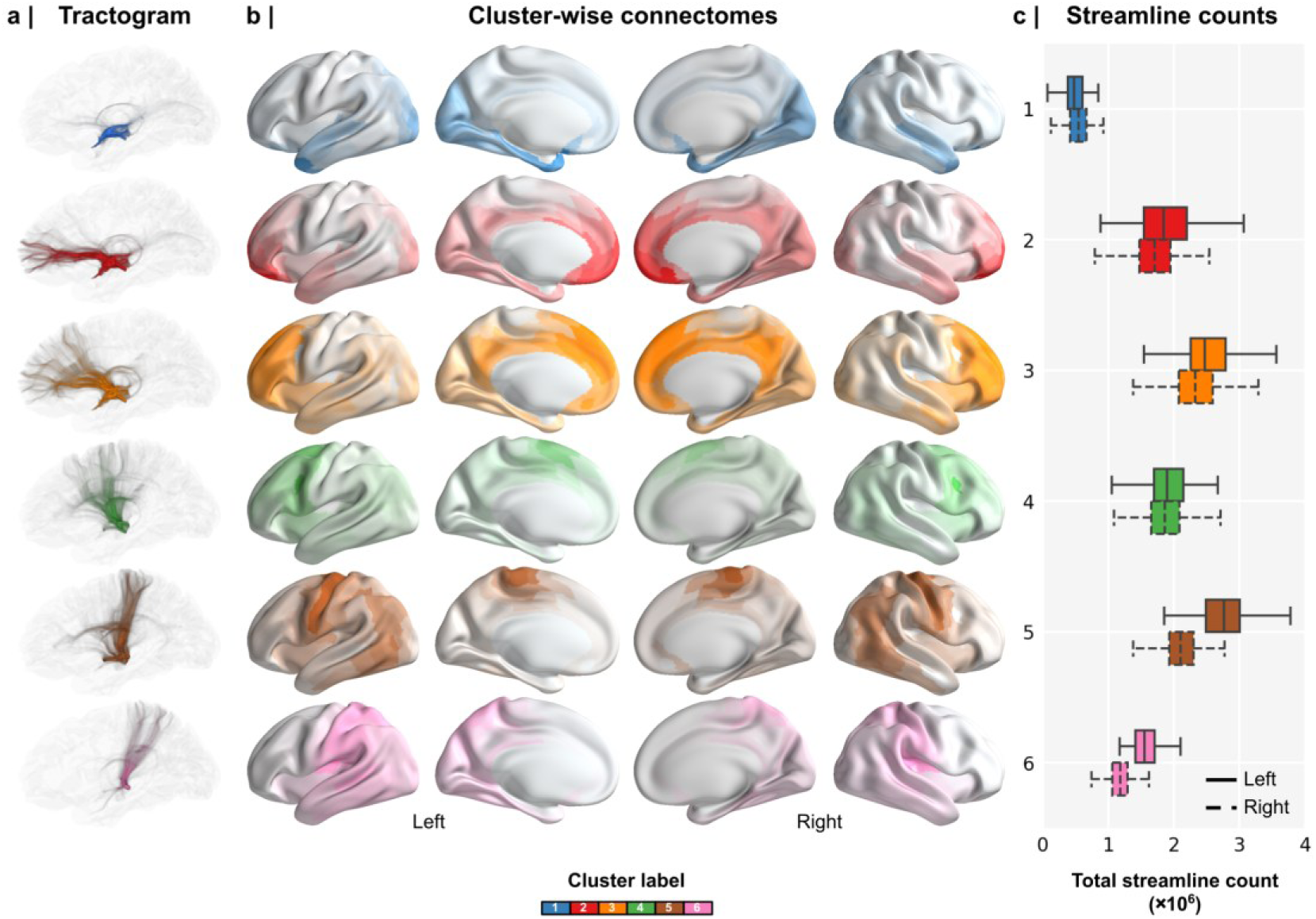
Cluster-wise tractography. (a) Example cluster-wise tractograms for the left hemisphere of a single subject. (b) Group-level cluster-wise cortical connectomes, where parcels are color-coded and have varying opacity levels (linearly scaled), reflecting the number of streamlines connecting each parcel with the clusters (row-wise). (c) Boxplot shows the total number of streamlines connecting the cortex with each cluster.

Using this analysis and similar to observations in rodent and non-human primate literature (Haber et al., 2023; Yang et al., 2022), we identified a central region of the ZI (Clusters 3-4) between the rostral (Clusters 1-2) and caudal (Cluster 5-6) regions that was demonstrated to be highly connected to the intervening dorsal prefrontal cortex (BA06, BA08) with decreasing frontopolar connections moving more caudally and increasing primary motor and sensory connections.

### Linking continuous and discrete connectivity findings

Having demonstrated the topographic organization of the cortico-incertal connectivity using diffusion map embedding as well as spectral clustering, how do the results using these two methods compare? To address this question, we used G1 and G2 to formalize a “2D gradient coordinate space” for the ZI, driven by the differences in cortical connectivity among individual voxels (Supplementary Figure 4a) and demonstrate the voxel-wise correspondence with the spectral clustering results. Combining G1 and G2 values (explaining 37% and 14% of the variance on average across left and right hemispheres, Supplementary Figure 2) in a 2D gradient coordinate space showed clear separation of the k=6 spectral clustering solution (Supplementary Figure 4b). This is reflected by an average Silhouette score of 0.324 (95% CI [0.311 – 0.337]) and 0.284 (95% CI [0.269 – 0.299]) for left and right hemispheres, which ranges from −1, indicating incorrect clustering, to 1, indicating highly dense clustering. Scores around zero indicate overlapping clusters. The 2D gradient coordinates were transferred to the ZI voxel and cortical surface space using a 2D colormap, and thus reduced in dimension to a single color code. This 1D color coding confirmed the dominant rostral-caudal axis among ZI voxels and their corresponding differences in connectivity from anterior prefrontal cortex to primary sensorimotor cortices (Supplementary Figures 4c and d).

### Tractographic evidence of a dorsal-ventral organization of the ZI

While our current results confirmed a predominant rostral-caudal topographic organization, evidence from animal models would suggest the presence of an additional dorsal-ventral organization (Mitrofanis, 2005; Monosov et al., 2022; Watson et al., 2014). Given the potential significance of this complementary organization, we specifically investigated coronally oriented reconstructions to assess for potential differences in cortical connectivity along this axis (Supplementary Figure 5) in comparison to the *in vivo* T_1_ maps (Lau et al., 2020), given these differences would not be easily detectable in the axial plane. Coronal cross-sections demonstrated high correspondence of G2 with the T_1_-based segmentation of the internal structures of the ZI seed label (in blue and magenta), based on the Schaltenbrand histological atlas (Lau et al., 2020; Morel et al., 1997; Schaltenbrand, 1977). A full range of coronal cross-sections is displayed in Supplementary Figure 6.

### Validating human cortico-incertal connectivity findings across datasets

Given that much of our analyses relied on high-resolution dMRI data obtained with optimized sequences on 7T MRI machines (Vu et al., 2015), which may not be widely accessible, it was important to assess the broader applicability of our findings. To ensure the technical validity of our results, we conducted several validation analyses. These included evaluating the replicability of our findings across images acquired with a lower magnetic field strength, assessing the reliability between test-retest sessions of the computational solutions, and confirming replicability at the individual subject level.

### Replicability between MRI field strengths and test-retest reliability

Visual inspection of the group-average gradients indicated overall high replicability and reliability across field strengths (3T and 7T) and the 3T test-retest datasets, respectively. The most notable differences were between 7T and 3T (Figure 4a-b). To quantify their similarity, Procrustes disparity was calculated for each dataset pair (where lower disparity indicates higher similarity). Quiver plots in Supplementary Figure 7 visually illustrate the shift in G1 and G2 values between datasets for each voxel in both the 2D coordinate as well as volume space. Quantitative analysis confirmed the highest similarity among 3T datasets (Figure 4c) versus those compared to the 7T dataset (average disparity score of 0.065 ± 0.016 vs. 0.014 ± 0.003 across both hemispheres, *p* < 0.001).

**Figure 4.**
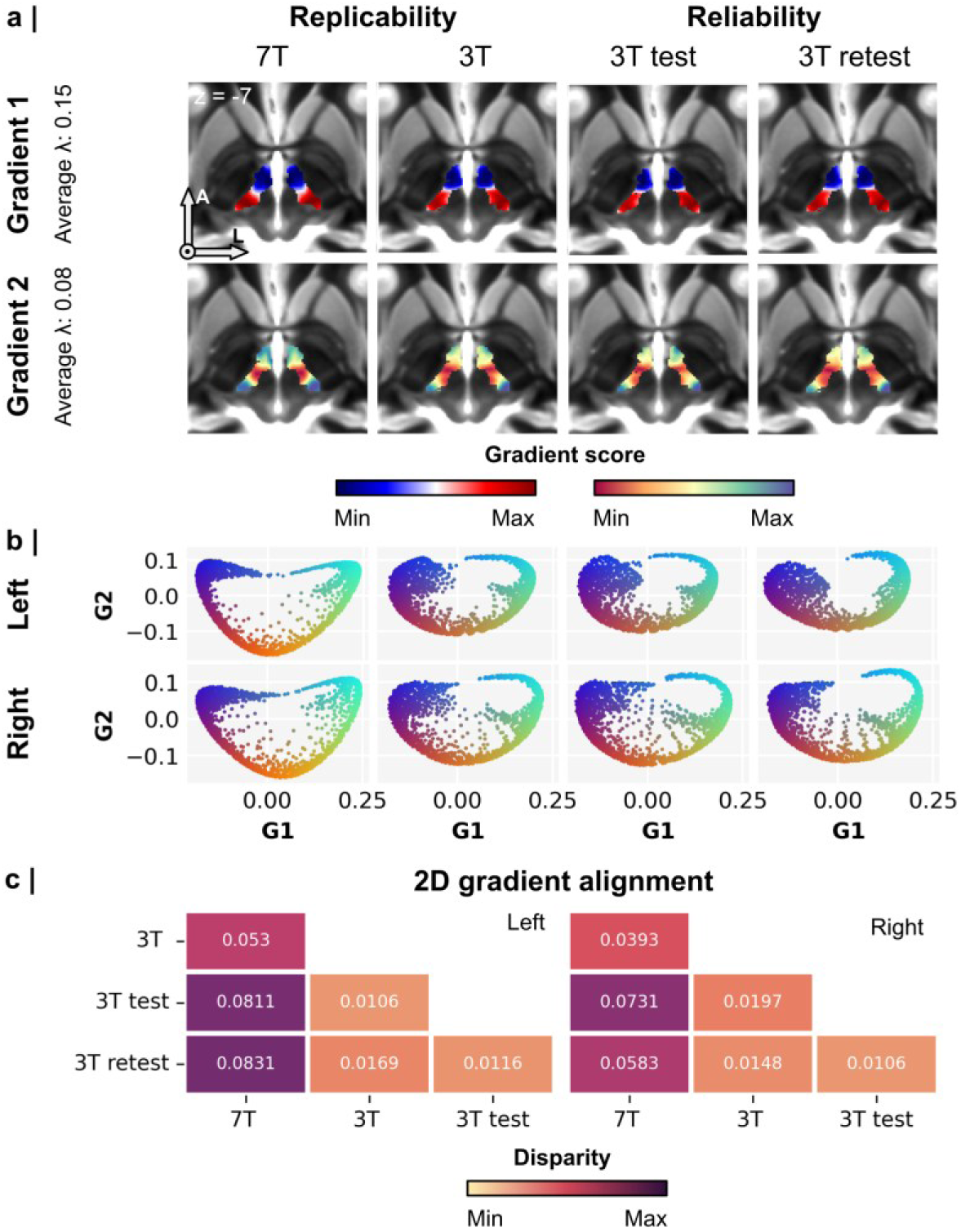
Replicability and reliability of zona incerta (ZI) connectivity patterns. (a) Axial cross-sections of gradient 1 and 2 per MRI dataset. (b) Similar to a but with ZI voxels displayed in the respective 2D gradient coordinates space. (c) Procrustes disparity scores for each comparison of 2D gradient coordinates among datasets.

Similar to the gradient data, the clustering results showed higher similarity among 3T datasets, as indicated by higher Dice coefficients and lower centroid distances (Figure 5). However, an average Dice score of 0.767 ± 0.042 and distance of 0.900 mm ± 0.199 mm between 7T and other datasets indicated good overlap or similarity. Notably, centroid distance results revealed a hemisphere-specific trend, with larger differences observed in the right hemisphere. Cluster-wise evaluation in Supplementary Figure 8 showed that in particular clusters 2 to 4 had the lowest replicability and reliability while clusters 5 and 6 had the highest.

**Figure 5.**
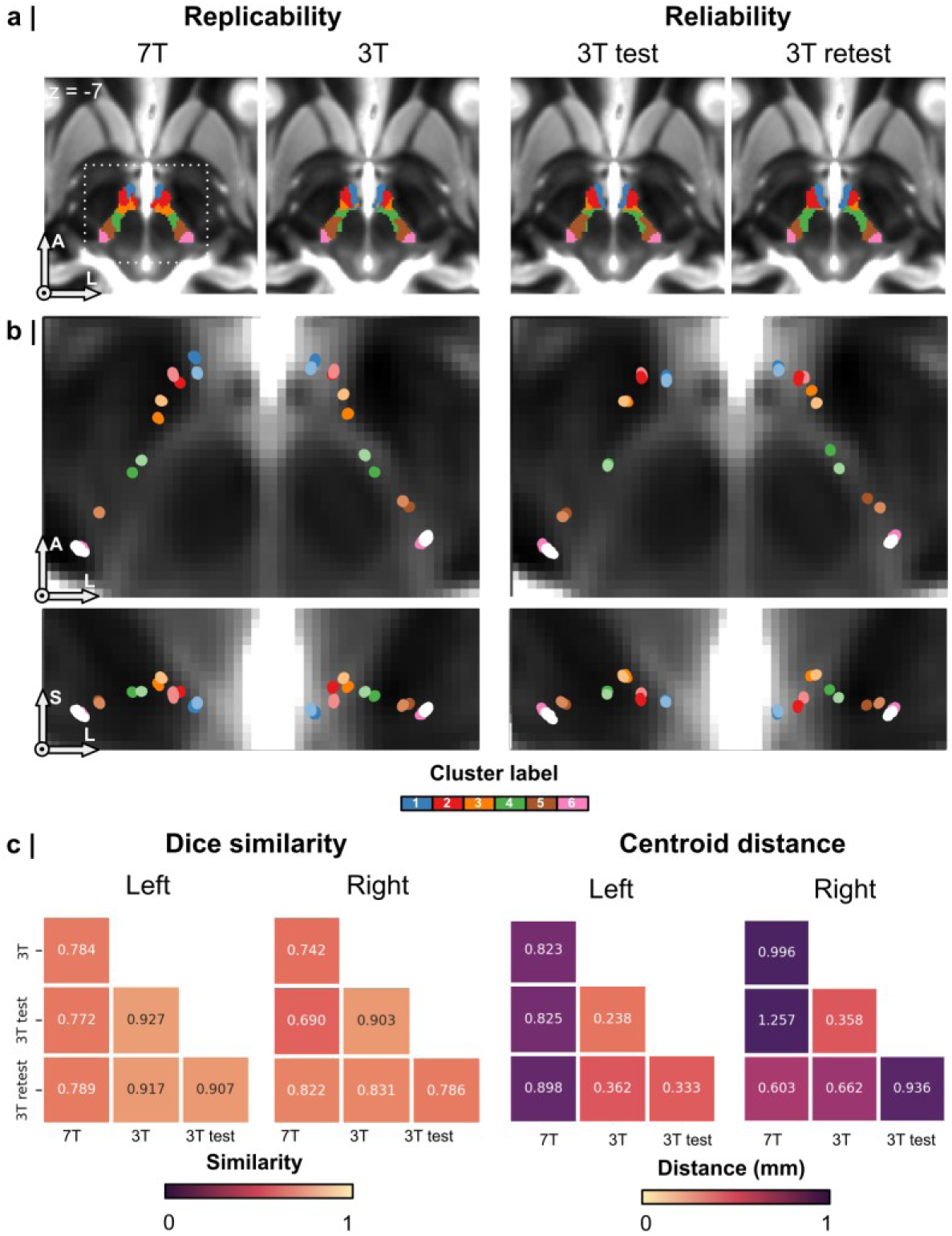
Replicability and reliability of the spectral clustering results. (a) Axial cross-sections of spectral clustering results (k=6) for each MRI dataset (left to right). (b) Comparison of cluster centroids between datasets, with results matched column-wise. Bright labels correspond to the 3T and 3T retest datasets. (c) Dice overlap scores and centroid distances for each comparison of spectral clustering results among datasets for the left and right hemispheres.

### Replicability at the individual subject level

Assessing the individual subject level replicability of the diffusion map embedding and clustering results helped to demonstrate the potential downstream utility of these techniques. High individual subject level replicability is a requisite for potential clinical applications such as DBS targeting. We found a high correlation between the group-based gradients and those from the individual subject’s data (mean Pearson’s correlation coefficient = 0.924 ± 0.052, Figure 6a). However, the primary gradient showed significantly higher values (0.963 ± 0.016) compared to the second gradient (0.886 ± 0.048, *p* < 0.001), consistent across both hemispheres. Compared to the between dataset comparisons (e.g., 3T vs. 7T), individual subject level cluster solutions relative to group-based clusters showed greater divergence, with centroid distances exceeding 1 mm (2.456 ± 0.246 mm) and Dice scores below 0.4 (0.228 ± 0.113, Figure 6b-c). Additionally, slightly larger differences were observed for the right hemisphere, as indicated by centroid distances, which varied among clusters more than Dice similarity.

**Figure 6.**
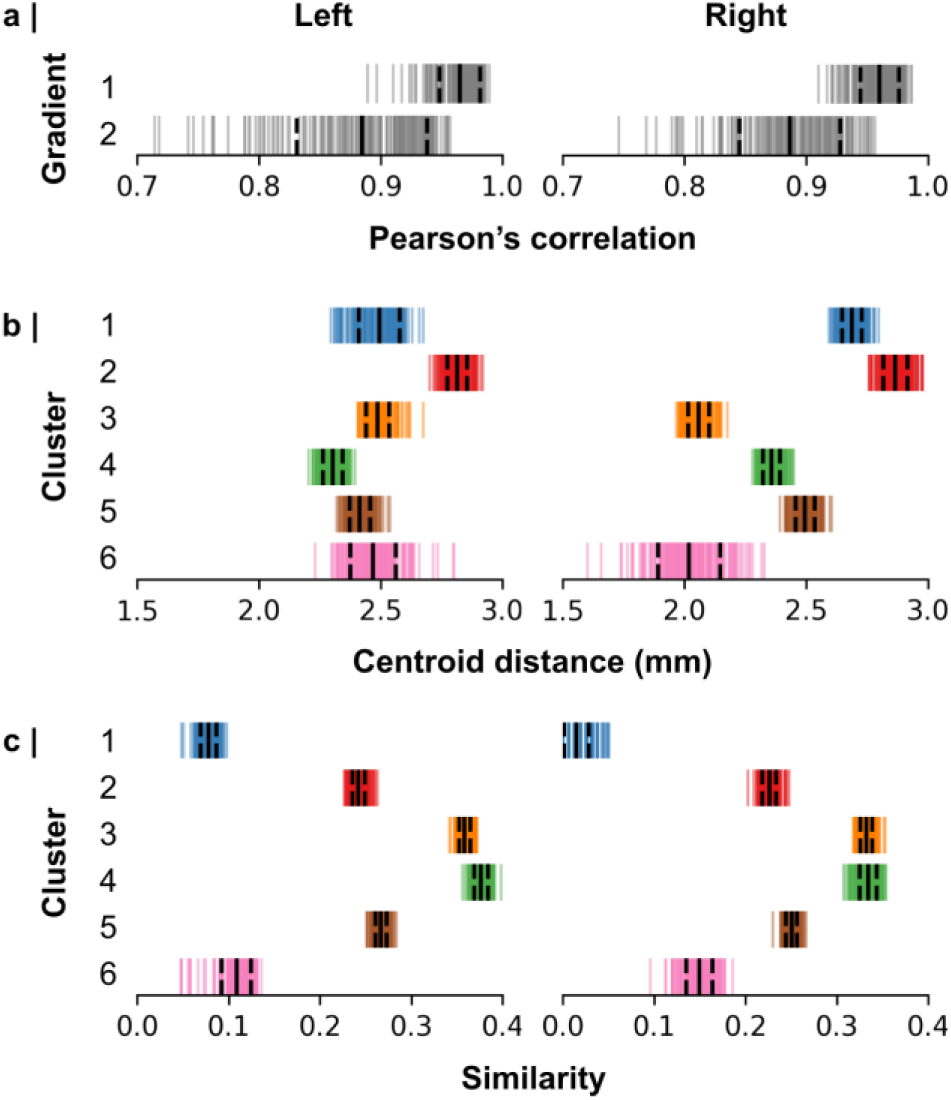
Individual subject replicability. (a) Spatial correlation between individual and group-level structural connectivity for each of the two retained gradients. (b) Centroid distances and (c) Dice scores for the alignment between individual and group-level clusters. Results are shown for both left and right hemispheres.

### Replicability using clinical-grade diffusion data

To evaluate the robustness and potential clinical applicability of our findings, we repeated the group-level analyses in an independently acquired diffusion MRI dataset (Kasa et al., 2022) using acquisition parameters that more closely resemble current clinical practice (2 mm isotropic resolution and a reduced number of diffusion-encoding directions). The principal rostro-caudal organization of cortico-incertal connectivity was preserved, indicating that the dominant topographic gradient is robust across imaging protocols and data quality (Supplementary Figure 9). In contrast, finer-grained parcellations showed reduced stability. While solutions with a small number of clusters remained broadly consistent with those obtained from the high-resolution HCP data, solutions with larger numbers (k > 4) exhibited increasing variability. These findings suggest that the large-scale organization of the human ZI can be recovered using clinically representative diffusion MRI, whereas reliable delineation of finer connectivity-defined subdivisions benefits from the higher spatial and angular resolution available in research-grade acquisitions (i.e. HCP and 7T).

### Linking cortico-incertal connectivity to cortical functions and hierarchies

After confirming the reliability of the structural connectivity estimates, we focused on understanding how the observed topographic organization relates to the functional role of the ZI. To achieve this, we compared the observed cortico-incertal connectivity patterns (Figure 2c) with known cortical maps of cognitive functioning and hierarchical organization, providing a context based on extent brain maps regarding the structural and functional implications of coricto-incertal connections. Examples of these cortical properties are shown in Supplementary Figure 10.

### Connectivity gradients

The cortical expression of G1 showed significant correlations with cognitive terms related to motor processing including “movement” (Pearson’s r = 0.680, *p* < 0.005), “coordination” (Pearson’s r = 0.565, *p* < 0.05), and “multisensory processing” (Pearson’s r = 0.562, *p* < 0.05, Figure 7a). G2 correlated strongly with terms such as “pain” (Pearson’s r = 0.479, *p* < 0.05), “response inhibition” (Pearson’s r = 0.444, *p* < 0.001), and “cognitive control” (Pearson’s r = 0.438, *p* < 0.001). Regarding cortical hierarchies (Figure 7b), G1 exhibited the strongest correlations with the first principal component of cognitive terms from the NeuroSynth database (“CogPC1”, Pearson’s r = 0.694, *p* < 0.005) (Yarkoni et al., 2011) and the magnetoencephalography theta-band activity (5–7 Hz) over the frontal regions (Pearson’s r = 0.821, *p* < 0.001) (Shafiei et al., 2022; Van Essen et al., 2012).

**Figure 7.**
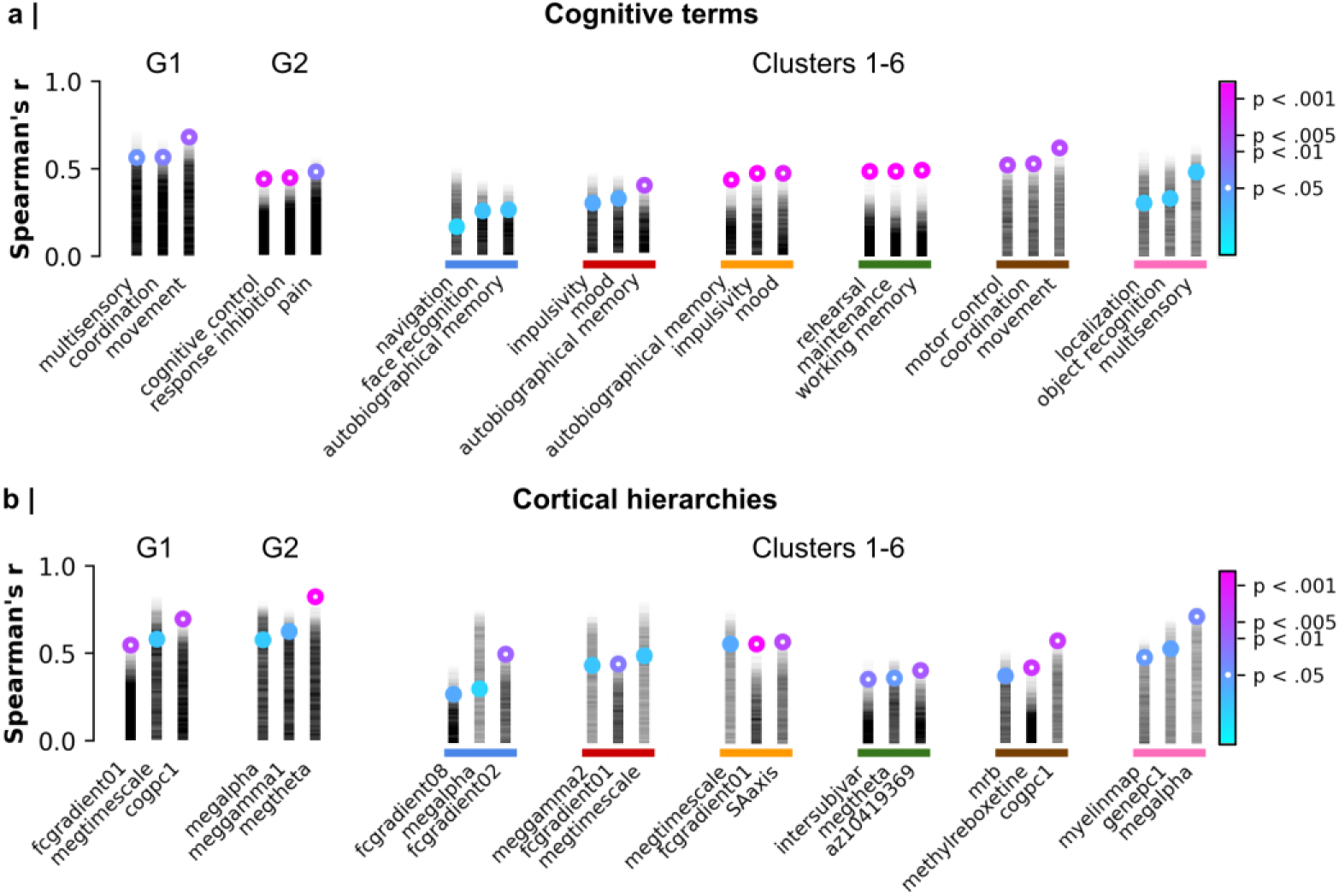
Correlation analysis between cortico-incertal structural connectivity patterns and cortical properties. The gradient-weighted cortical maps (Figure 2c) and cluster-wise connectomes (Figure 3) exhibit significant correlations with various (a) cognitive terms (e.g., movement) and (b) cortical hierarchies (e.g., sensorimotor-association axis) based on Spearman’s correlation coefficient (colored markers). Marker colors represent P-values, corrected for spatial autocorrelation using N=10k spin tests. Semi-transparent black bars show permuted values. The source file containing all Spearman’s correlation coefficients is available in the code repository referenced in the manuscript.

### Spectral clustering

Strong correlations with cognitive terms and cortical hierarchies were particularly evident for clusters 3-5 (Figure 7a-b), which have the highest number of connecting streamlines (Figure 3c). Cluster 5, located near the central sulcus, is significantly linked to movement-related cognitive processing (Pearson’s r = 0.623, *p* < 0.005) and CogPC1 (Pearson’s r = 0.593, *p* < 0.005). Cluster 4, situated more anteriorly, overlaps with regions involved in working memory (Pearson’s r = 0.494, *p* < 0.001) with a high level of expression of serotonin (5-HT1B) receptors (Pearson’s r = 0.413, *p* < 0.005) (Beliveau et al., 2017). Cluster 3, located further towards the frontal pole, is associated with mood (Pearson’s r = 0.473, *p* < 0.005) and impulsivity (Pearson’s r = 0.473, *p* < 0.001), and the sensorimotor-association axis (Pearson’s r = 0.583, *p* < 0.005) (Sydnor et al., 2021). Clusters 1, 2 and 6, characterized by the least number of connecting streamlines (Figure 3c), were relatively weakly associated (i.e., low Spearman’s coefficient and/or within the spatial autocorrelation range) with cognitive terms or cortical hierarchies.

### Clinical relevance of cortico-incertal connectivity for stereotactic targeting

Finally, to evaluate the implications of data-driven ZI structural connectivity characterization for stereotactic targeting, we assessed the overlap of the stimulation volume obtained in a patient with essential tremor (ET) treated successfully using cZI DBS, with ZI voxels in their 2D gradient coordinate space and clusters (Figure 8a-b). This patient notably had complete resolution of tremor symptoms based on the essential tremor rating assessment scale at one year follow-up post-DBS with absence of side effects, and thus could aid in defining optimal stimulation targets based on the cortico-incertal structural connectivity. For the left hemisphere, ZI voxels within the stimulation volume were characterized by mean G1 and G2 scores of 0.066 ± 0.058 and −0.130 ± 0.036, respectively, and overlapped predominantly with cluster 4 (∼64%, Figure 8c). The right hemisphere stimulation value was characterized by mean G1 and G2 scores of 0.174 ± 0.045 and −0.046 ± 0.061, respectively, and overlapped with clusters 4 (∼22%) and 5 (∼20%).

**Figure 8.**
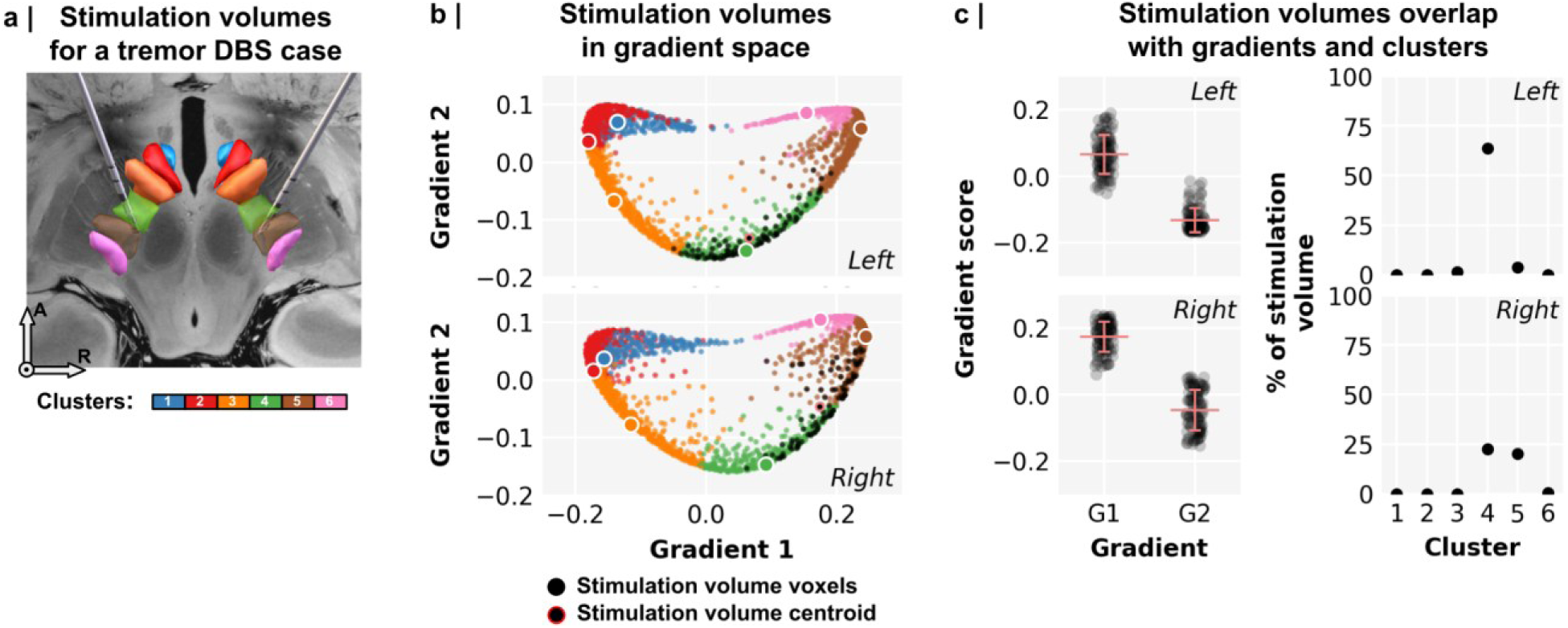
Mapping a deep brain stimulation (DBS) case electrode stimulation volume to ZI tractography-based gradients and clusters. (a) DBS electrode reconstruction was performed using the Lead-DBS (v3.0.0) software and visualized in a common space overlayed with the ZI clusters derived in this work. (b) Propagated coordinates of the left and right stimulation volumes (black markers) into gradient space. Stimulation volume centroids are shown with red outlines. (c) Distribution of gradient scores within stimulation volumes and percent (%) of the DBS stimulation volume overlapping with ZI clusters.

## Discussion

Traditionally understood as “a space between structures”, our understanding of the ZI has evolved to recognize distinct subregions connected with the neocortex, which provide the anatomical basis for location-dependent functional variability (Chometton et al., 2017; Z. Li et al., 2021; Lin et al., 2023; Mitrofanis et al., 2004; Moriya and Kuwaki, 2020). In this study, we investigated the cortico-incertal connections in humans through probabilistic tractography, reporting differential connectivity patterns and their functional implications within the ZI. One key finding stemming from this approach is the identification of a topographic organization that follows a rostral-caudal axis, with evidence for a central ZI region situated between the better characterized rostral and caudal regions of the human ZI. We demonstrate that gradient and spectral clustering approaches provide complementary strategies for elucidating the connectivity of substructures within the ZI. Combined with information extracted from contextual analyses, these findings offer valuable insight regarding the structural organization of the ZI and how it relates to function and targeted therapy.

### Cortico-incertal connectivity follows a topographic organization in humans

The human ZI exhibits extensive connections throughout the neocortex with predominant connectivity to the frontal lobe, followed by the parietal lobe, and relatively restricted connectivity with the temporal and occipital lobes (Supplementary Figure 1; Supplementary Table 1). Probabilistic tractography further revealed a topographic organization of the cortico-incertal connections along the rostral-caudal axis (Figure 2), validated by replicability and reliability analyses from both the 3T and 7T datasets (Figures 4 and 5). Specifically, we observed that the rZI exhibits robust connectivity with the anterior and ventral PFC as well as the rostral and subgenual ACC. Conversely, the cZI demonstrates prominent connectivity with the primary motor, premotor, and somatosensory cortices. These findings mirror evidence from previous tract tracing studies in rodents (Mitrofanis and Mikuletic, 1999; Yang et al., 2022) and non-human primates (Coudé et al., 2018; Haber et al., 2023; Ongür et al., 1998; Saluja et al., 2024), demonstrating separable connectivity patterns between rZI and cZI subregions. Specifically, rodent studies show the rZI receives inputs from the frontal lobe and the prelimbic areas, while the cZI receives inputs from the motor areas and the basal ganglia (Yang et al., 2022). Non-human primate studies revealed a similar topographic pattern, with the rZI receiving projections from the prefrontal cortex and the cZI connecting to motor areas (Haber et al., 2023; Ongür et al., 1998).

### The primary gradient follows a rostral-caudal pattern of organization

A rostral-caudal cortico-incertal connectivity pattern is directly visible in the primary gradient (G1, Figure 2). Specifically, the rZI exhibits extensive streamlines anchored to the prefrontal cortex, including the anterior prefrontal cortex (BA10) and the anterior cingulate cortex, whereas the cZI is preferentially anchored with the primary sensory and motor cortices (Figure 2). These results are in line with previous tractography and axon tracing experiments (Haber et al., 2023; Saluja et al., 2024). The rostral and caudal ZI were connected to associated cortical regions by way of the anterior and posterior limbs of the internal capsule, respectively providing a structural substrate for the functional diversity observed across ZI subregions, which at the cortical level are significantly associated with coordination, sensory integration, and movement as determined via contextual analyses (Figure 7). These results are also in line with theories regarding the rostral-caudal organization of the granular PFC, from the perspective of a hierarchy of cognition that becomes increasingly more complex more rostrally, before integrating with emotional control from the agranular ventromedial and orbitofrontal cortices (Badre and D’Esposito, 2009; Koechlin and Summerfield, 2007; Levy, 2024). Previous studies in mice report that inputs from the prefrontal region to the rZI are associated with regulating escape speed (Chou et al., 2018; Hormigo et al., 2023), curiosity (Ahmadlou et al., 2021; Monosov et al., 2022), fear (Z. Li et al., 2021; Venkataraman et al., 2021; L. Zhang et al., 2022; Zhou et al., 2021, 2018), and anxiety processing (Z. Li et al., 2021; L. Zhang et al., 2022; Zhou et al., 2021). Consistent with rodent literature, recent primate studies provide evidence that the rZI acts as an integrative hub that connects cortical and subcortical targets modulating fast, survival-based responses in a dynamic environment (Haber et al., 2023). Similarly, previous findings also indicate a connection between the sensorimotor cortex and cZI, highlighting its role in modulating motor control. Selective activation of cZI neurons has been shown to alleviate motor symptoms in Parkinsonian mice (L. X. Li et al., 2021). Tract tracing experiments have further elucidated projections from the motor cortex to the ZI in non-human primates (Haber et al., 2023; May and Basso, 2018; Saluja et al., 2024). Taken together, the primary gradient reported in this study aligns with current knowledge of the rostral and caudal ZI’s involvement in diverse neurological processes, highlighting the distinct connectivity patterns of these ZI subregions to their respective cortical regions as the structural basis for their functional roles.

### The secondary gradient reveals a central region in the human ZI

A second gradient (G2) was identified that is anchored on connections between the central ZI and the dorsal PFC separable from cortico-incertal connections with the rostral and caudal ZI respectively (Figure 2). To our knowledge, this is the first report of separable cortical connections to the central ZI region in humans. Supporting these results, tract tracing experiments in rodents have reported the central sectors exhibit distinct cortical connectivity patterns compared to the rostral and caudal regions (Yang et al., 2022). Detailed macaque tract tracing work has also identified clear connections between the ACC and dorsal PFC and the central ZI both dorsally and ventrally (Haber et al., 2023). G2 was found to be significantly associated with terms such as “pain”, “cognitive control”, and “response inhibition”, suggesting an integrative role of the ZI in pain processing and decision making. In line with this, optogenetic and chemogenetic experiments demonstrate that selective activation of the ZI reduces pain perception in mice (Hu et al., 2019; Li et al., 2023, 2022; Moriya and Kuwaki, 2020; Wu et al., 2023) and rats (Moon et al., 2016), whereas selective inhibition increases pain perception (Moon et al., 2016; Moriya et al., 2020; Wu et al., 2023). DBS of the ZI has been demonstrated to alleviate thermal-related pain in PD patients (Lu et al., 2021), suggesting a role for pain-related modulation.

The pattern of G2 aligns with previous findings of high resolution longitudinal T_1_ mapping in the rZI (Lau et al., 2020), as voxels with higher G2 values correspond to longer T_1_ relaxation time (Supplementary Figure 5). While this appears to follow a dorsal-ventral organization that has been captured in rodents (Mitrofanis, 2005; Paxinos and Watson, 2006; Watson et al., 2014) and non-human primate models (FitzGibbon et al., 2000; Haber et al., 2023; Hardman and Ashwell, 2012; May and Basso, 2018; Mitrofanis et al., 2004), it more likely separates the rostral subregion from the surrounding fields of Forel (Lau et al., 2020). This aligns with our previous demonstration that T_1_ mapping serves as a robust method for visualization of the ZI *in vivo* at 7T, separating the rostral and caudal subregions from surrounding white matter tracts (Lau et al., 2020). Building on this work, we report that probabilistic tractography permits detailed visualization of the ZI in humans, identifying a consistent organization of cortico-incertal connections.

### Discrete subregions of the zona incerta using spectral clustering

While diffusion map embedding uncovered smooth transitions of cortical connections within the ZI (Margulies et al., 2016; Royer et al., 2024), mapping discrete subregions of the ZI to discrete cortical regions is a conventional, and furthermore intuitive, approach as well (Petersen et al., 2024). To explore this alternative view, we assessed the topographic organization of the ZI using spectral clustering, identifying boundaries between clusters of voxels with similar connectivity patterns. Cluster-wise findings are potentially more relatable to tract-tracing literature (Donahue et al., 2016) and may offer higher certainty compared to voxel-wise analysis, which try to resolve features at a sub-areal scale (Petersen et al., 2024).

Inspired by the cytoarchitectonic classification of the rodent ZI into six sectors (Romanowski et al., 1985), we employed a spectral clustering solution (k=6) to account for this known diversity, which furthermore provides a level of granularity for investigating location-dependent neuromodulation along the ZI (Awad et al., 2020; Bingham and McIntyre, 2024; Blomstedt et al., 2018; Mostofi et al., 2019; Plaha et al., 2011, 2006; Shepherd et al., 2024). The resulting clusters followed the same rostral-caudal topographic organization observed with diffusion map embedding (Figure 2, Figure 3). Our spectral clustering results show that rostral clusters (Clusters 1-2) are preferentially connected to cortical areas located more at the frontopolar and ventral prefrontal cortex, thus associated with cognitive and emotional processing (e.g., “facial recognition,” “mood,” “impulsivity,” “autobiographical memory”). The centrally located Clusters 3-4 connect with dorsal PFC regions, contextually linked to memory-related terms (“working memory,” “maintenance,” “rehearsal”), while caudal clusters (Clusters 5-6) connected with premotor, and M1 and S1 are associated with sensorimotor functions (“locomotion,” “coordination,” “multi-sensory”). This spectral clustering approach was found to provide evidence complementary to the gradient approach for revealing connections between premotor areas and a central ZI region, separable from the rostral ZI, which connects to the prefrontal cortex, and the cZI, which connects to the primary motor region. While the association between the rZI with cognition (Haber et al., 2023; Z. Li et al., 2021; Lin et al., 2023) and the cZI with locomotion are well established (Awad et al., 2020; L. X. Li et al., 2021), our contextual analysis reveals that the central ZI regions are implicated in prefrontally-mediated memory processing in humans. Supporting this, the ZI has been shown in mice to be essential for memory formation employing methods involving selective inhibition (Sun et al., 2024; Zhou et al., 2018). Please note that these contextual findings should be viewed as hypothesis-generating for future work designed to directly link ZI subregions with behavior, cognition, and neuromodulation, rather than definitive evidence of functional specialization.

### Enhancing stereotactic targeting of the ZI through topographic mapping

Mapping the connectivity profile along the rostral-caudal axis of the ZI can help elucidate its integrative role as a neuromodulation target for a wide spectrum of neurological disorders (Awad et al., 2020; Bingham and McIntyre, 2024; Blomstedt et al., 2018; Mostofi et al., 2019; Plaha et al., 2011, 2006; Shepherd et al., 2024). The structural connectivity gradients and cluster solutions derived in this study, provide a reference framework to project DBS electrode coordinates (Figure 8). We demonstrate how these topographic maps can be employed for visualizing stimulation sites in the context of cortico-incertal connections. Compared to spectral clustering, the presented 2D gradient coordinate space provides a unique visualization of ZI organization that could be employed for refining targeting, optimizing the therapeutic window, and complementing recent developments in this area of research to identify DBS sweet spots (Hollunder et al., 2024).

### Challenges of *in vivo* tractography and recommendations

Integrating dMRI findings with validation methods such as tract tracing from the literature is crucial for assessing the accuracy of connectivity studies (Thomas et al., 2014). Our observations align well with previous tract tracing evidence from the rodent or non-human primate studies, supporting the reliability of dMRI and tractography in exploring cortico-subcortical connectivity reliably (Kai et al., 2022). However, it is important to acknowledge the spatial limitations of anatomical fidelity in these techniques (Maffei et al., 2022; Maier-Hein et al., 2017).

Identifying fine tracts, such as the dorsal-ventral organization within the ZI, is constrained by the quality of dMRI data, including spatial resolution and signal-to-noise-ratio (SNR). To address these challenges, we used the 7T HCP dataset, which was acquired with MRI protocols optimized for high spatial resolution (1.05 mm^3^) and SNR dMRI data (Vu et al., 2015). Furthermore, by focussing primarily on larger (cortico-incertal) white matter tracts, the inherent risk of spurious tracts is mitigated, and our validation analyses support this, showing replicability at lower magnetic field strengths, acquisition protocols, and at the level of individual subjects (Figures 4-6). Note that the individual replicability metrics are sensitive to the quality of the spatial transformations between image spaces (e.g., native vs. template). Integration of highly sensitive metrics of image coregistration quality, for example, by the placement of anatomical fiducials on structural MRI scans, can be considered in future work (“A framework for evaluating correspondence between brain images using anatomical fiducials,” n.d.; Taha et al., 2023). Alternatively, individualized ROIs can be considered to prevent the need for coregistrations between different image spaces. Moreover, the lower individual replicability observed for the spectral clustering results (i.e., relative to the gradients) are likely due to the stronger impact of single voxel differences on the respective evaluation metrics.

Tractography relies on assumptions regarding water diffusion directionality in tissues that may potentially lead to inaccuracies, particularly when dealing with complex fiber orientations as observed in subcortical regions (Axer et al., 2011; Basser and Pierpaoli, 1996). Each step in the processing workflow involves decisions that impact subsequent steps and results (Jeurissen et al., 2019; F. Zhang et al., 2022). For example, mask selection influences the number of streamlines identified; larger masks yield more streamlines, while smaller masks may miss connections. Mask accuracy is crucial, as overlap with gray matter or cerebrospinal fluid can cause anatomically implausible connections. Our ZI mask was based on extensive intra- and inter-rater reliability assessments limiting the potential for inaccuracy in labeling (Lau et al., 2020). The choice of tractography algorithm is another important consideration. Deterministic algorithms may result in more false-negatives (missed connections), while probabilistic algorithms may produce more false-positives (implausible connections). Additionally, the parameters for defining tractography termination criteria must balance sensitivity and specificity.

Finally, given the limitations of *in vivo* tractography, several aspects of our findings warrant further investigation. First, strong evidence supports connectivity between the ZI and subcortical structures (Mitrofanis, 2005). Since our analyses focused only on cortico-incertal connections, future work should also examine the role of subcortical pathways (Kai et al., 2022). Second, the dorsal-ventral organization reported in rodent and non-human primate brains may relate to the connectivity patterns observed in humans. However, current evidence (e.g., variability in G2) remains inconclusive and requires targeted follow-up. Third, although we performed several validation analyses, including an independently acquired diffusion MRI dataset with clinically representative acquisition parameters (2 mm isotropic resolution and a reduced number of diffusion-encoding directions), further work is needed to establish the minimum data quality requirements for robust connectivity-based ZI parcellation. While the principal rostro-caudal organization was preserved across acquisition protocols, finer connectivity-defined subdivisions were less reproducible with the lower-resolution data, suggesting that characterization of subtle topographic features continues to benefit from the higher spatial and angular resolution afforded by research-grade diffusion MRI.

### Concluding remarks

Using computational neuroanatomy approaches grounded in probabilistic tractography, we aimed to improve the understanding of the topographic organization of the human ZI and its connections with the cerebral cortex. Our findings highlight a distinct rostral-caudal gradient that links specific cortical areas to corresponding regions within the ZI. This detailed mapping, validated across multiple datasets and at the individual subject level, supports the involvement of the ZI across a diverse range of functions, from motor control to cognitive control and emotional regulation. The complementary evidence provided by gradient and spectral clustering approaches enhance our understanding of the structural organization of the ZI with potential applications for optimizing targeted neuromodulation strategies.

## Materials and methods

### Datasets

#### High-quality MRI data

Minimally pre-processed data as part of the Human Connectome Project (HCP) young adults study (Glasser et al., 2013; Van Essen et al., 2012) were used to evaluate the structural connectivity of the ZI through probabilistic tractography. Structural connectivity analyses using the dMRI data were carried out on three separate datasets: “7T” (i.e., the reference), “3T” and “3T test-retest” (N=36; 11M/25F, aged 22-35). All subjects included in the 7T dataset (N=169, 68M/101F; aged 22-35) were included in the 3T dataset (N=169, 68M/101F; aged 22-35).

For each subject, T_1_-weighted (T_1_w) 3T MRI scans were acquired with a 3D MPRAGE sequence (Mugler III and Brookeman, 1990): resolution = 0.7 mm isotropic voxels; repetition time/echo time (TR/TE) = 2400 / 2.14 ms. Acquisition of dMRI data varied between field strengths. The 7T dMRI images were collected on a Siemens MAGNETOM 7T MRI system (Vu et al., 2015) with a 1.05 mm3 nominal isotropic voxel size, TR=7000 ms, TE=71.2 ms, b-values=1000, 2000 s/mm2 (64 directions per shell), FOV=210 × 210 mm2 with 15 b-value = 0 s/mm^2^ images. For the 3T and 3T test-retest datasets, dMRI data were collected with a 1.25 mm^3^ nominal isotropic voxel size, TR=5520 ms, TE=89.50 ms, b-values=1000, 2000, 3000 s/mm^2^ (90 directions per shell), FOV=210 × 180 mm^2^ with 18 b-value = 0 s/mm^2^ images. The 3T and 3T test-retest datasets were acquired on customized Siemens Skyra 3T MRI systems. Full acquisition details are described in the HCP1200 reference manual^1^.

#### Clinical-quality MRI data

An additional MRI dataset (N=23, 10/14 M/F, mean age = 36±15 years) acquired with a clinical-grade protocol was obtained using a 3T MRI system (Siemens Prisma, Erlangen Germany) with a 32-channel head coil. The scanning protocol included the acquisition of structural images using the MPRAGE sequence: TR/TE=5000/2.98 ms, TI=700 ms, FOV=256 × 256mm2, 1 mm^3^ isotropic voxel size. The dMRI data were collected with a 2 mm^3^ nominal isotropic voxel size, TR=2800 ms, TE=66.80 ms, b-values=1300, 2600 s/mm^2^ (130 diffusion-encoding directions acquired twice with left–right, right-left phase encoding directions), FOV=224 × 224 mm^2^ with 20 b-value=0 s/mm^2^ images. These data from control participants were acquired as part of a previous study (Kasa et al., 2022).

### Structural connectivity

#### Probabilistic tractography

Probabilistic tractography was performed to derive a structural connectivity matrix from the dMRI data. As part of the minimal preprocessing pipeline data release (Glasser et al., 2013), all subjects underwent FreeSurfer processing (v5.3.0-HCP) (Fischl, 2012). The ZI ROI mask was first resampled and transformed to the individual subjects’ minimally preprocessed volume space (0.7mm^3^) (Lau et al., 2020). Volumetric neocortical labels were built by mapping the HCP-MMP1.0 surface parcellation using Connectome Workbench’s ribbon-constrained *label-to-volume-mapping* function and FreeSurfer-derived surfaces (Glasser et al., 2016; Marcus et al., 2011). The ZI mask (thresholded at 50% and dilated with a 3 voxel radius) was used to seed tractography to the target neocortical regions using FSL’s *probtrackx* with 5000 streamlines per ZI ROI voxel (Behrens et al., 2007). The resulting probability maps in the ZI quantified the number of streamlines that reached each target. Generated connectivity was then transformed from subject’s native space to the MNI152NLin6Asym template space and reduced to a 2-dimensional *M*-by-*N* matrix, where *M* represents the voxels in the ZI ROI (1981 and 1901 for left and right hemispheres, respectively) and *N* is the neocortical targets (180 each hemisphere) with their corresponding number of streamlines.

#### Connectivity gradients

Nonlinear dimension reduction using diffusion map embedding was used to transform the connectivity matrices into a low-dimensional representation (Coifman et al., 2005). ZI voxels that are characterized by similar connectivity patterns will have a value closer together in the low-dimensional space, whereas voxels with little or no similarity are farther apart. A total of hundred connectivity gradients were calculated using the *GradientMaps* function in the BrainSpace Python toolbox (Vos de Wael et al., 2020). These were computed in two ways: using (i) the group-averaged and (ii) the individual subject’s (for individual subject level replicability analyses, see respective section) connectivity matrices as input, while using the normalized angle kernel and the diffusion map embedding approach, and otherwise default parameter settings (Vos de Wael et al., 2018).

#### Spectral clustering

In parallel to the gradients, spectral clustering of the structural connectivity matrices was performed using the *SpectralClustering* function in the scikit-learn library for Python, using the default parameters and a chosen solution (i.e., number of clusters) ranging from k=2 to k=8 (Pedregosa et al., 2011). As for the calculation of gradients, spectral clustering calculates an affinity matrix from the connectivity matrices and performs a low-dimension embedding but adds an additional K-means clustering to assign discrete labels to each voxel. As a result, voxels belonging to the same cluster have similar connectivity patterns.

#### Neocortical connectivity maps

We generated neocortical connectivity maps of the spectral clustering and gradients results to visualize and evaluate the ZI connectivity in terms of its relation to the neocortex and its properties. First, gradient-weighted neocortical maps were created by multiplying each row of the ZI-neocortical connectivity matrix with the corresponding gradient value of that ZI voxel (Guell et al., 2020; Katsumi et al., 2023). Second, the cortex was associated with the ZI spectral clustering result by assigning each target neocortical region with the label of the cluster demonstrating the greatest connectivity (i.e., a “winner-takes-all” based on the maximal number of streamlines).

Finally, cluster-wise neocortical connectivity maps were extracted, based on the average number of streamlines connecting the voxels within each cluster label to each neocortical parcel for contextual analysis.

### Validation analyses

#### 7T vs. 3T replicability and test-retest reliability

##### Connectivity gradients

To enable comparison between datasets (e.g., 7T vs. 3T), all connectivity gradients (N*=*100) were aligned to the 7T dataset using Procrustes shape analyses. Procrustus analysis performs scaling/dilation, rotations, and reflections of the input gradients to match the reference 7T gradients and minimize the sum of the squares of the pointwise differences (i.e., disparity) between the two input datasets, *A* and *B*:

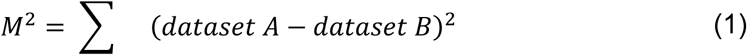

This is particularly useful in the case of comparison of *N*D gradient spaces, which are characterized by arbitrary scalings. The resulting, aligned gradients were then mapped back onto the ZI voxel space to visualize and further analyze structural connectivity patterns. The same analysis was used to quantify the similarity of the ZI structural connectivity patterns among datasets based on the first two gradients (i.e., 2D gradient space) by calculating their disparity scores.

##### Spectral clustering

For the evaluation of the spectral clustering results, we used the Dice coefficient and centroid distance for each cluster. For the Dice coefficient, voxels of the cluster are first binarized before comparing their overlap by identifying the ratio of corresponding non-zero voxels across the two datasets to the total number of non-zero voxels across both datasets:

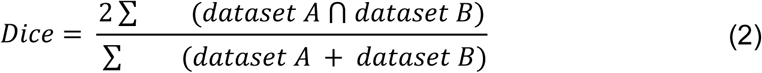

To calculate the centroid distance, the coordinates of the centroid for each cluster is first determined before a Euclidean distance is calculated between corresponding pairs of centroids across datasets.

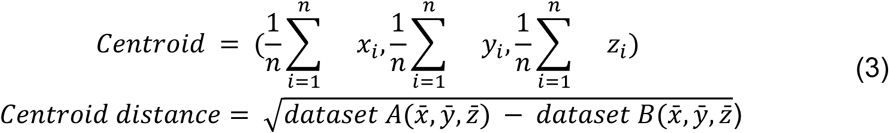

Together, these two measures offer insights into the similarities of corresponding labels across the datasets. A higher Dice coefficient is better and indicative of greater volumetric overlap. The Dice coefficient is typically interpreted as follows: poor (<0.2), fair (0.2 – 0.4), moderate (0.4 – 0.6), good (0.6 – 0.8) and excellent (>0.8) overlap (Kreilkamp et al., 2019). On the other hand, a shorter centroid distance is preferable, suggesting that the extent of similarity of volumes are minimal. It can also provide insight into the extent at which the volumes differ along the spatial components.

##### Individual subject level replicability

In addition, considering the potential application of these gradients and spectral resulting results for stereotactic targeting at the individual level, individual subject level replicability in the 7T dataset was assessed as well. We estimated the spatial correlation between individual and group-level structural connectivity for each of the two retained gradients (Yang et al., 2020). Similarly, we also calculated Dice scores and centroid distances to further evaluate the alignment between individual subject and group-level clusters (“A framework for evaluating correspondence between brain images using anatomical fiducials,” n.d.; Kai et al., 2022).

##### Contextual analyses

The gradients-based and cluster-wise neocortical connectivity maps were compared with brain-wide cognitive terms and cortical hierarchies to evaluate the functional and biological relevance of the ZI structural connectivity.

First, we used the NeuroSynth meta-analytic database (neurosynth.org) to assess how the neocortical connectivity maps of the ZI gradients and cluster-specific connectomes relate to cognitive terms (Yarkoni et al., 2011). NeuroSynth aggregates data from over 14,000 published functional MRI studies to derive probabilistic measures of the association between brain voxels and cognitive processes based on frequently occurring keywords such as “pain” and “attention”. Similar to the approach in Hansen (2023) based on the Cognitive Atlas, a public ontology that provides an extensive list of neurocognitive processes (Poldrack et al., 2011), we utilized 124 cognitive and behavioral terms, covering broad categories (“attention”, “emotion”), specific cognitive processes (“visual attention”, “episodic memory”), behaviors (“eating”, “sleep”), and emotional states (“fear”, “anxiety”).

Second, we used the neuromaps toolbox to assess whether the ZI neocortical connectivity maps are linked to molecular, microstructural, electrophysiological, developmental, and other functional properties (N=73) of the neocortex (Markello et al., 2022). See “Statistical analysis” for more details.

##### Clinical relevance

Finally, as a proof-of-concept, we reconstructed the stimulation volumes from a single DBS case targeting the ZI to characterize the structural connectivity patterns of the voxels located within the stimulation volumes in both hemispheres.

##### Data acquisition

Pre-operative T_1_-weighted (T_1_w) and T_2_-weighted (T_2_w) 7T MRI scans were acquired. T_1_w imaging was performed using the 3D MP2RAGE pulse sequence (Marques et al., 2010): resolution = 0.8 mm isotropic voxels; repetition time/echo time (TR/TE) = 6000/2.69 ms; inversion times = 800/2700 ms; flip angles = 4/5°. T_2_w-imaging was performed using a 2D SPACE pulse sequence with parameters: resolution = 0.6 mm isotropic voxels and TR/TE = 2000/131 ms. During preprocessing, images were corrected for gradient non-linearity distortions. MP2RAGE data were additionally corrected for B_1_ ^+^-inhomogeneities using a separately acquired Sa2RAGE B_1_ ^+^ map (1.9×1.9×2.8 mm voxels; TR/TE = 2400/0.81 ms; TI1/TI2 = 45/1800 ms; flip angles = 4/11°) (Eggenschwiler et al., 2012; Haast et al., 2018; Marques et al., 2010).

##### Stimulation volume reconstruction

We employed Lead-DBS 3.0 (lead-dbs.org) (Horn and Kühn, 2015) for electrode localization and reconstruction in a common space (see Figure 8a). In brief, a linear co-registration was performed between the pre-operative, preprocessed T_1_w MRI and post-operative CT. Subsequently, the 7T T_1_w and T_2_w images were non-linearly registered to the ICBM 2009b Nonlinear Asymmetric (i.e., MNI2009bAsym) (Fonov et al., 2009) template space. All registrations were performed in accordance with previously validated presets (Avants et al., 2008) of the Advanced Normalization Tools (ANTs; <u>stnava.github.io/ANTs/</u>) (Avants et al., 2011) as implemented in Lead-DBS. The SimBio algorithm (Oostenveld et al., 2011; Wolters et al., 2006) was used to estimate the stimulated volume based on stimulation parameters at least 1-month after the DBS device had been turned on. The essential tremor rating assessment scale scores were time-locked with DBS stimulation parameters.

## Statistical analyses

For each cortical map derived from the NeuroSynth and neuromaps databases (see “Contextual analyses” section), we parcellated it using the HCP-MMP1.0 atlas and computed its Pearson’s spatial correlation with the ZI neocortical connectivity maps, accounting for spatial autocorrelations using N=10,000 spin tests. During each spin test, the parcel coordinates were randomly rotated, and original parcels were reassigned the value of the closest rotated parcel according to the Hungarian algorithm (Kuhn, 1955). Additionally statistical evaluations were performed using the appropriate tests implemented in the *pingouin* v0.5.3 Python package.

## Data and code availability

The non-clinical data used in this study are available as part of the publicly available Human Connectome Project S1200 release (humanconnectome.org/study/hcp-young-adult). Anonymized stimulated volumes and clinical scores are available at github.com/ataha24/zona-clusters. The raw clinical data are available upon request. Analysis was performed using a combination of publicly available toolboxes: Connectome Workbench v2.0 (Marcus et al., 2011), BrainSpace v0.1.10 (Vos de Wael et al., 2020), NeuroSynth v0.3.8 (Yarkoni et al., 2011), neuromaps v0.0.3 (Markello et al., 2022), BrainStat v0.3.2 (Larivière et al., 2023), and custom Python code available at the code repository. A full list of installed Python packages is listed in the repository as well.

## Acknowledgements

Data were provided in part by the Human Connectome Project, WU-Minn Consortium (Principal Investigators: David Van Essen and Kamil Ugurbil; 1U54MH091657) funded by the 16 NIH Institutes and Centers that support the NIH Blueprint for Neuroscience Research; and by the McDonnell Center for Systems Neuroscience at Washington University. Author RAMH was supported by a H2020 Marie Skłodowska Curie Actions Postdoctoral Fellowship (101061988). Author ARK was supported by the Canada Research Chairs program #950-231964, NSERC Discovery Grants RGPIN-2015-06639 and RGPIN-2023-05558, Canadian Institutes for Health Research Project grant #366062, Canada Foundation for Innovation (CFI) John R. Evans Leaders Fund project #37427, the Canada First Research Excellence Fund, and Brain Canada. Author JCL was supported by an NSERC Discovery Grant RGPIN-2023-05562 and research start-up funding from the Department of Clinical Neurological Sciences at Western University.

## Supplementary Figures

**Supplementary Figure 1.**
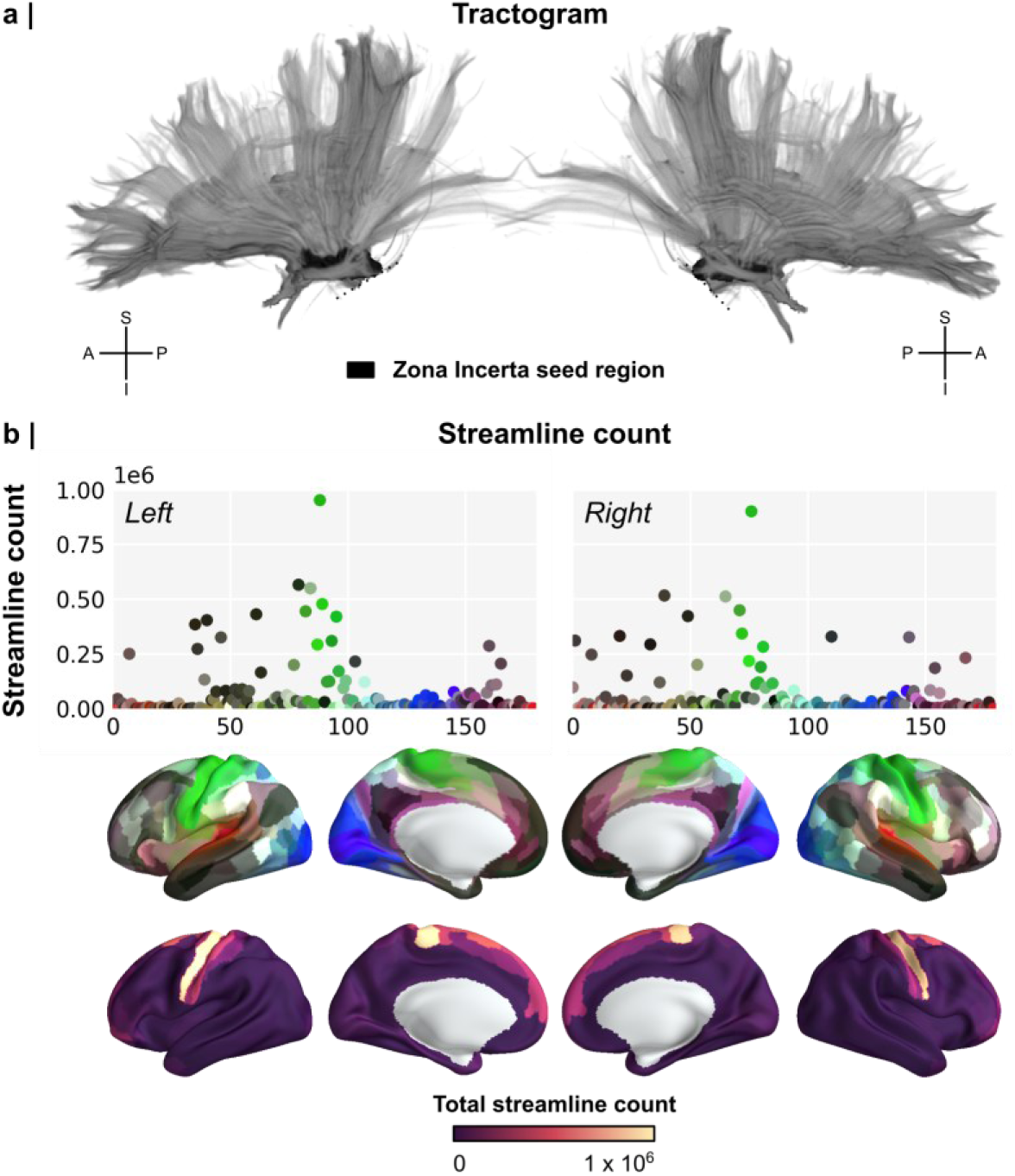
Spectral clustering tractograms and streamline count. (a) Tractogram based on the 7T MRI dataset shown from left vs. right views. (b) Scatter plots showing the streamline count per cortical parcel, color-coded according to the HCP-MMP1.0 atlas as well as a cortical surface representation, color-coded according to the number of streamlines. The source file containing streamline counts is available in the code repository referenced in the manuscript.

**Supplementary Figure 2.**
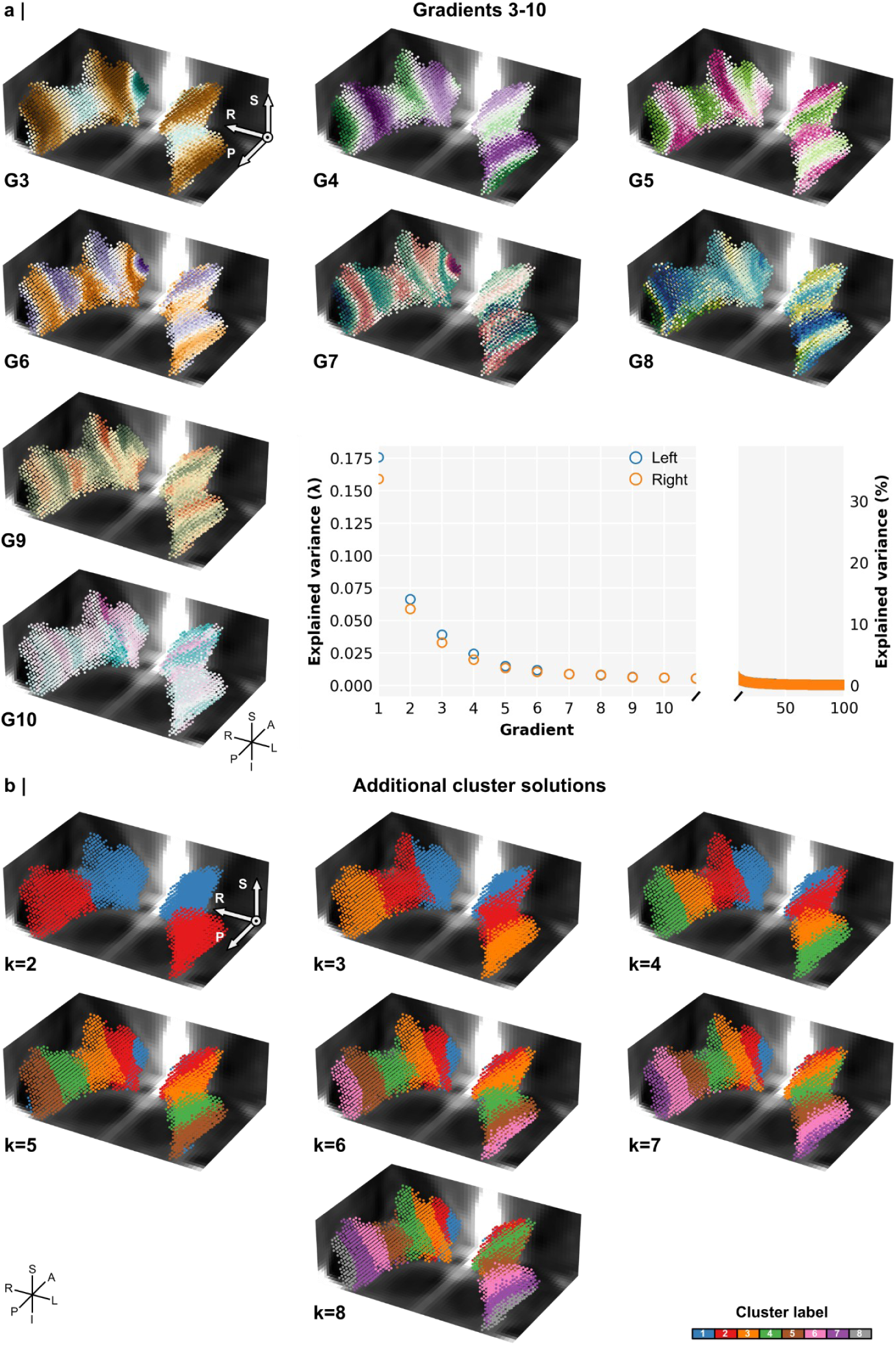
Overview of successive gradients and alternative cluster solutions. (a) Gradients 3-10 (G3-10) shown using 3D radiological views. The scree plot shows the variance explained by each gradient as indicated by their lambda value (left y-axis). The right y-axis displays each gradient’s respective contribution (%) to the total explained variance by all gradients. (b) Other spectral clustering solutions (k=2-8).

**Supplementary Figure 3.**
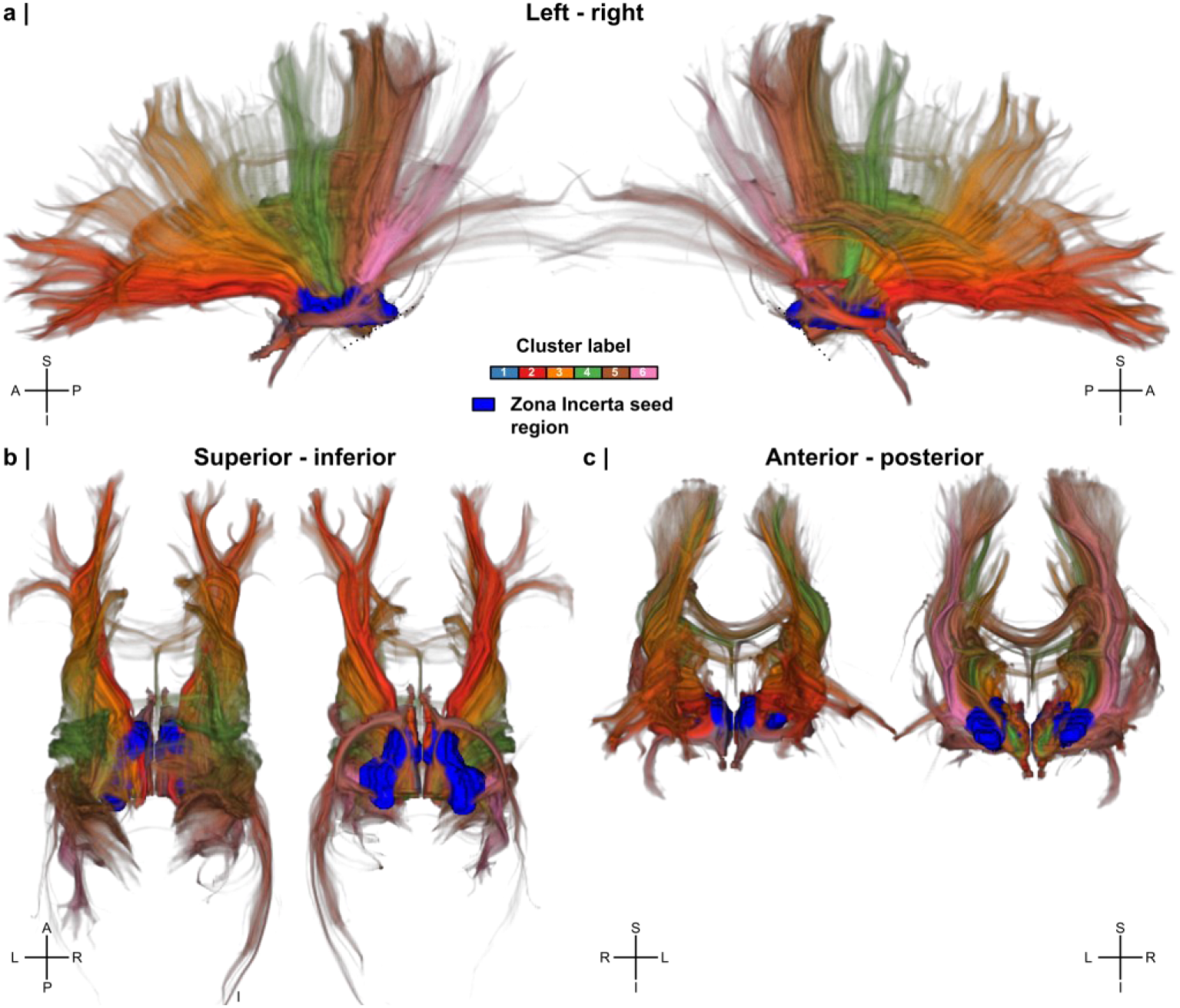
Composite cluster-wise tractogram. (a) Composite tractogram colored by spectral clustering labels, based on the 7T MRI dataset and for k=6 clusters solution, shown fom left vs. right, (b) superior vs. inferior and (c) anterior vs. posterior views.

**Supplementary Figure 4.**
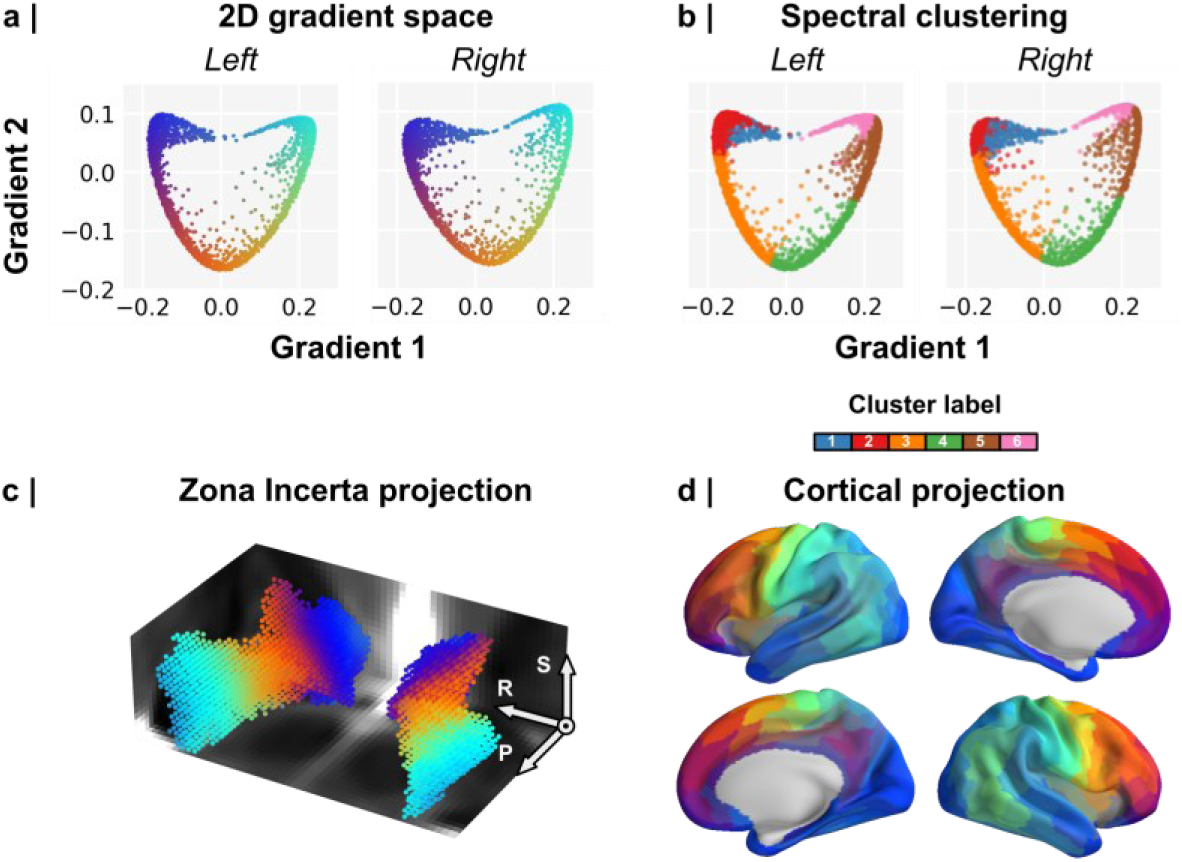
Zona incerta (ZI) 2D gradient coordinate space. (a) Gradient 1 (x-axis) and 2 (y-axis) values were used to position each ZI voxel in the corresponding 2D gradients coordinates space, color-coded using a 2D colormap. (b) Similarly as a, but ZI voxels are color-coded for the k=6 spectral clustering solution. (c-d) Projection of the 2D gradient coordinate space onto the ZI volume and cortical surface spaces, respectively.

**Supplementary Figure 5.**
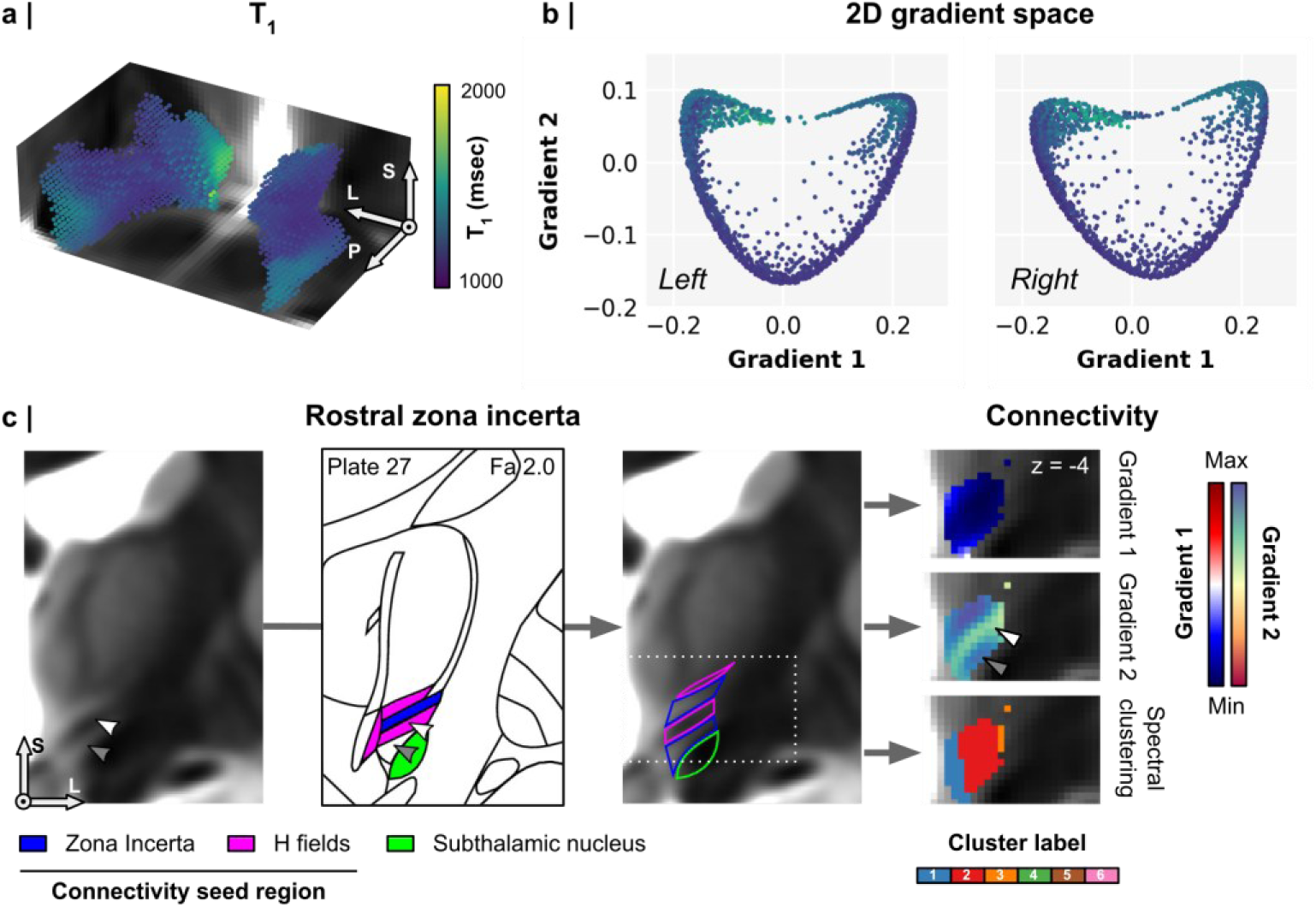
Zona incerta (ZI) longitudinal relaxation times (T_1_). (a) 3D volumetric, radiological display of ZI T_1_ values (msec). (b) ZI voxels in the 2D gradient coordinate space color-coded for T_1_ value. Voxels with high gradient 2 values are characterized by longer T_1_ values. (c) Comparison of the rostral ZI T_1_, Schaltenbrand atlas, and gradient and spectral clustering results (left to right). Rostral ZI T_1_ differences align with changes in gradient 2 values.

**Supplementary Figure 6.**
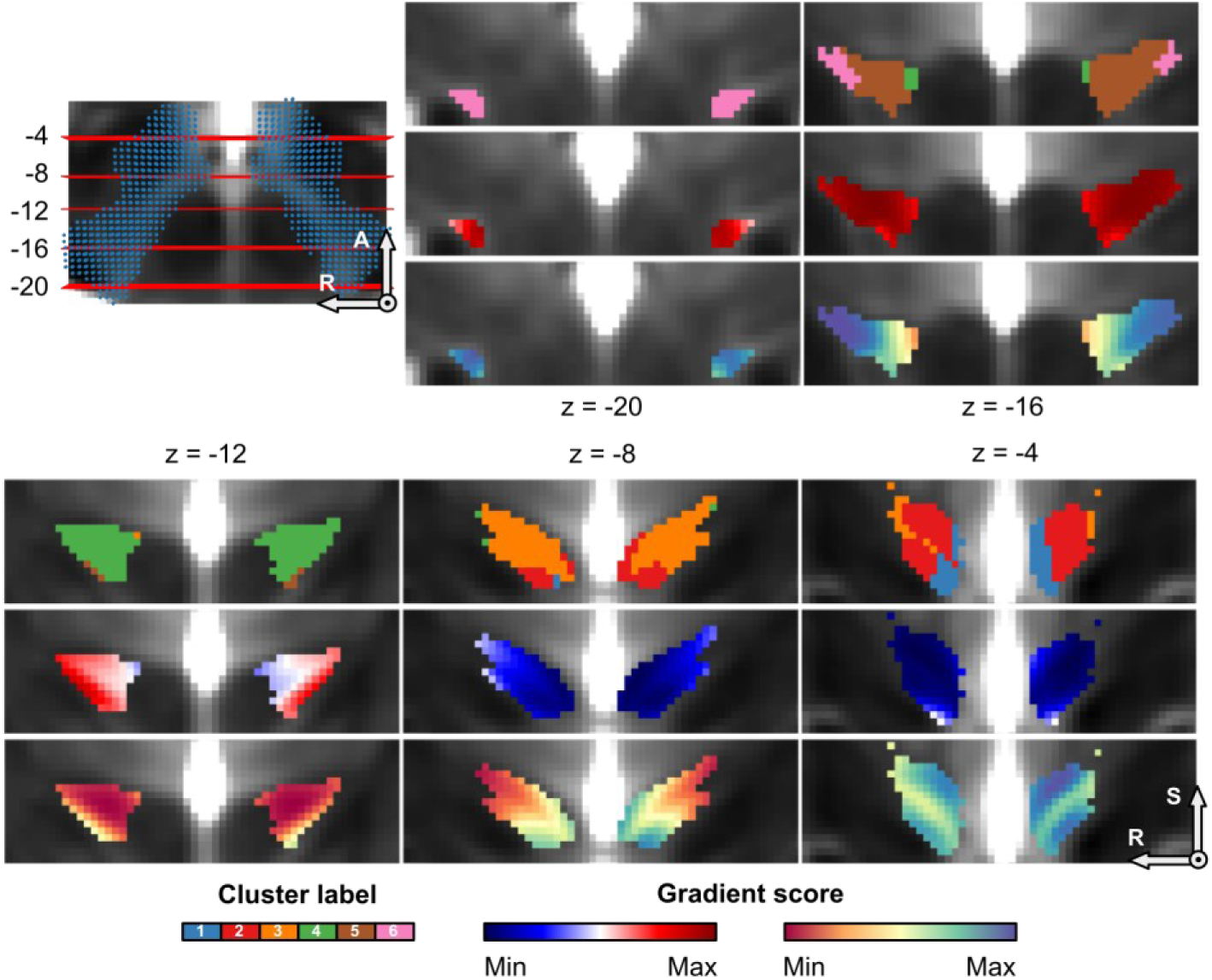
Coronal cross-sections of zona incerta gradient and spectral clustering results.

**Supplementary Figure 7.**
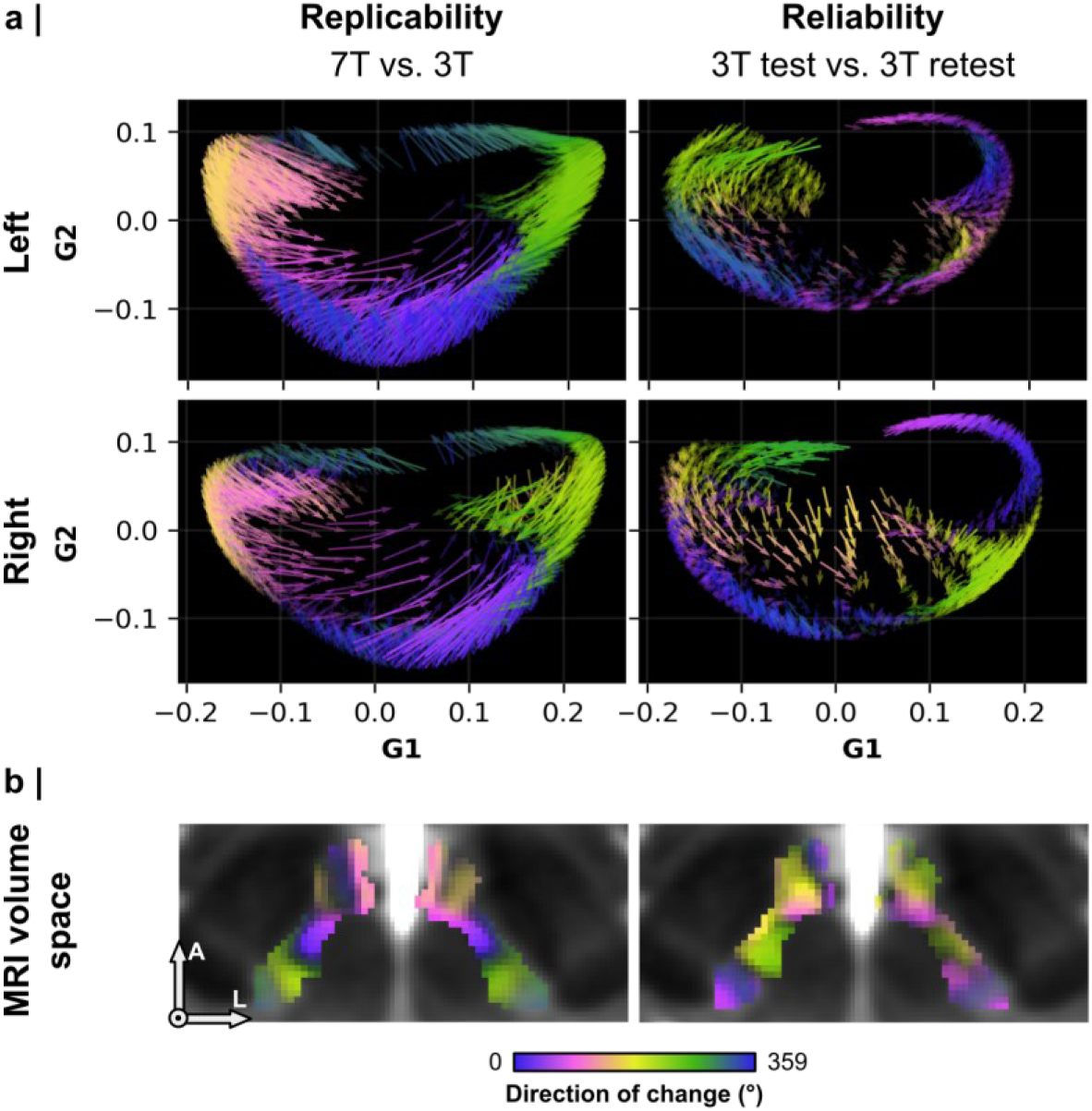
Visual display of the comparison between MRI datasets in the 2D gradient coordinate space. (a) Each arrow in the quiver plots illustrate the shift in gradient 1 (G1) and gradient 2 (G2) values between datasets for a single voxel, color-coded for the direction of change. (b) Projection of the quiver plots into the MRI volume space for localization purposes.

**Supplementary Figure 8.**
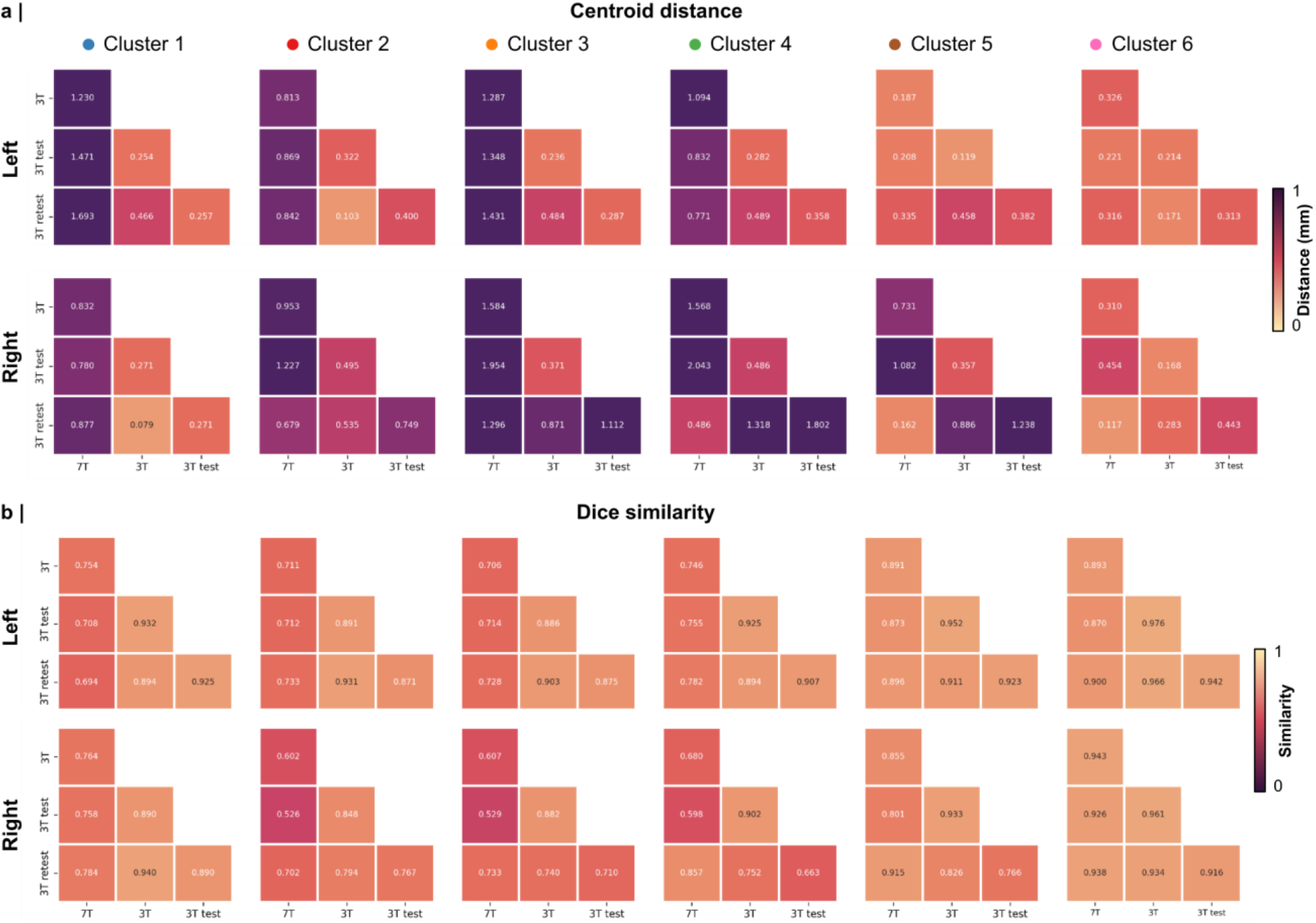
Replicability and reliability of the spectral clustering results. (a) Centroid distance (mm) and (b) Dice similarity scores, split per hemisphere and cluster.

**Supplementary Figure 9.**
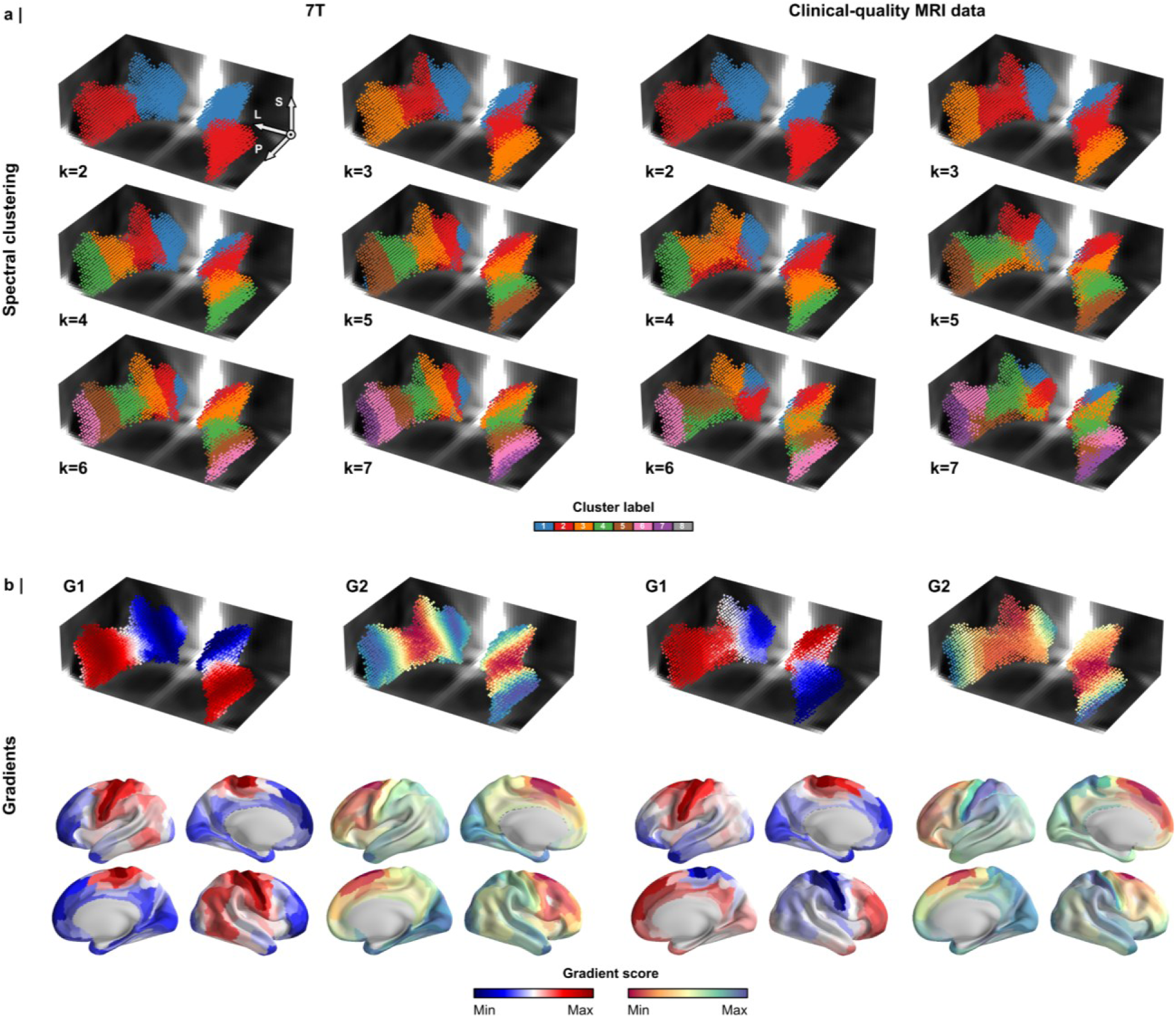
Comparison of spectral clustering and connectivity gradients across diffusion MRI acquisition protocols. (a) Spectral clustering results and (b) connectivity gradients obtained from the high-resolution 7T HCP dataset and an independently acquired clinical-quality diffusion MRI dataset (2 mm isotropic resolution with a reduced number of diffusion-encoding directions).

**Supplementary Figure 10.**
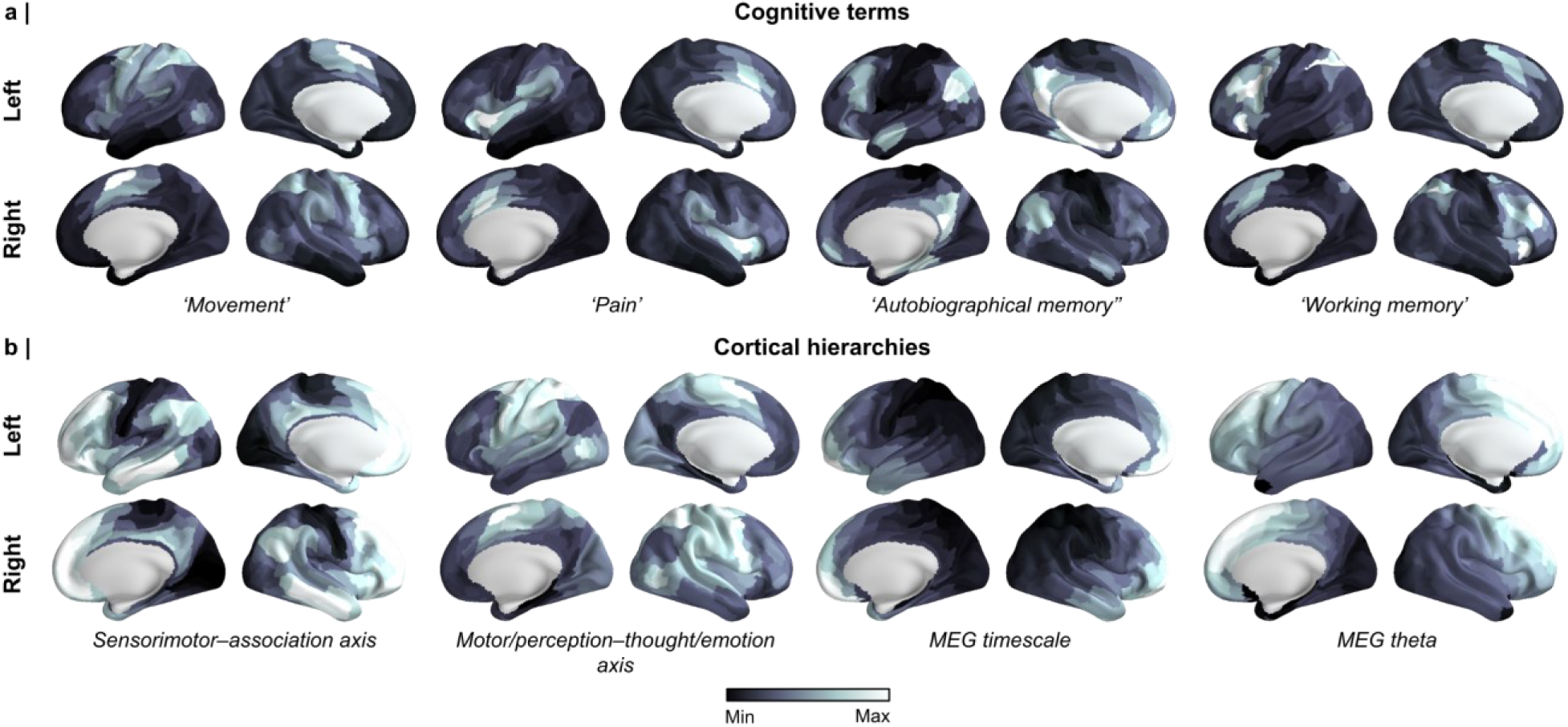
Example cortical (a) NeuroSynth (N=124) and (b) neuromaps (N=73) maps used for contextual analysis. Each cortical map is scaled according to their minimum and maximum value.

---

**Supplementary Table 1.** Top 5% of connections between each zona incerta (ZI) cluster (k=6) and cortical regions. Cortical regions from the left and right hemispheres (HCP-MMP1.0) with streamline counts exceeding the 95^th^ percentile for each cluster are ranked in descending order. The source files used for generating these rankings are available in the code repository referenced in the manuscript.

|  |  | Cluster 1 |  | Cluster 2 |  | Cluster 3 |  | Cluster 4 |  | Cluster 5 |  | Cluster 6 |  |
| --- | --- | --- | --- | --- | --- | --- | --- | --- | --- | --- | --- | --- | --- |
|  |  | HCP-MMP | Brodmann | HCP-MMP | Brodmann | HCP-MMP | Brodmann | HCP-MMP | Brodmann | HCP-MMP | Brodmann | HCP-MMP | Brodmann |
| Left | 1 | PeEc | Temporal | 10d | BA10 | 8BL | BA08 | 6ma | BA06 | 4 | BA04 | 1 | BA01 |
|  | 2 | TGd | Temporal | a10p | BA10 | SFL | BA06 | SFL | BA06 | 6mp | BA06 | 3b | BA03 |
|  | 3 | 10d | BA10 | a47r | BA47 | 9m | BA09 | 6mp | BA06 | 3b | BA03 | 4 | BA04 |
|  | 4 | Pir | Temporal | 10pp | BA10 | 8BM | BA08 | 4 | BA04 | 1 | BA01 | 2 | BA02 |
|  | 5 | a10p | BA10 | 9m | BA09 | 10d | BA10 | SCEF | BA06 | 3a | BA03 | 3a | BA03 |
|  | 6 | V1 | V1 | p10p | BA10 | 9a | BA09 | 6d | BA06 | 6d | BA06 | 7AL | BA07 |
|  | 7 | PreS | Temporal | 9a | BA09 | a47r | BA47 | 8BL | BA08 | 6ma | BA06 | 7Am | BA07 |
|  | 8 | a47r | BA47 | 11l | BA11 | 6ma | BA06 | 6a | BA06 | 5m | BA06 | 5L | BA05 |
|  | 9 | 10pp | BA10 | 8BL | BA08 | 9p | BA09 | 8BM | BA08 | 2 | BA02 | 5m | BA05 |
|  |  | Cluster 1 |  | Cluster 2 |  | Cluster 3 |  | Cluster 4 |  | Cluster 5 |  | Cluster 6 |  |
|  |  | HCP-MMP | Brodmann | HCP-MMP | Brodmann | HCP-MMP | Brodmann | HCP-MMP | Brodmann | HCP-MMP | Brodmann | HCP-MMP | Brodmann |
| Right | 1 | 10pp | BA10 | 10pp | BA10 | 8BL | BA08 | 6ma | BA06 | 4 | BA04 | 4 | BA04 |
|  | 2 | a10p | BA10 | a10p | BA10 | SFL | BA06 | SFL | BA06 | 6mp | BA06 | 3b | BA03 |
|  | 3 | V1 | V1 | 10d | BA10 | 9m | BA09 | 6mp | BA06 | 3b | BA03 | 1 | BA01 |
|  | 4 | TGd | Temporal | 9a | BA09 | 9a | BA09 | 4 | BA04 | 3a | BA03 | 3a | BA03 |
|  | 5 | PeEc | Temporal | 9m | BA09 | 8BM | BA08 | SCEF | BA06 | 1 | BA01 | 2 | BA02 |
|  | 6 | 10d | BA10 | a47r | BA47 | 6ma | BA06 | 8BL | BA08 | 6ma | BA06 | 5L | BA05 |
|  | 7 | 8BL | BA08 | p10p | BA10 | 10d | BA10 | 6d | BA06 | 6d | BA06 | 7AL | BA07 |
|  | 8 | 9m | BA09 | 8BL | BA08 | a47r | BA47 | 8BM | BA08 | SFL | BA06 | 5m | BA05 |
|  | 9 | 9a | BA09 | 11l | BA11 | a10p | BA10 | s6-8 | BA06/BA08 | SCEF | BA06 | 7Am | BA07 |

## Footnotes

1 https://www.humanconnectome.org/study/hcp-young-adult/document/1200-subjects-data-release

## References

A framework for evaluating correspondence between brain images using anatomical fiducials. n.d. DOI: 10.1002/hbm.24693

Ahmadlou M, Houba JHW, van Vierbergen JFM, Giannouli M, Gimenez G-A, van Weeghel C, Darbanfouladi M, Shirazi MY, Dziubek J, Kacem M, de Winter F, Heimel JA. 2021. A cell type-specific cortico-subcortical brain circuit for investigatory and novelty-seeking behavior. Science (New York, N.Y.) 372:eabe9681. DOI: 10.1126/science.abe9681, PMID: 33986154

Arena G, Londei F, Ceccarelli F, Ferrucci L, Borra E, Genovesio A. 2024. Disentangling the identity of the zona incerta: a review of the known connections and latest implications. Ageing Research Reviews 93:102140. DOI: 10.1016/j.arr.2023.102140

Avants BB, Epstein CL, Grossman M, Gee JC. 2008. Symmetric diffeomorphic image registration with cross-correlation: Evaluating automated labeling of elderly and neurodegenerative brain. Medical Image Analysis 12:26–41. DOI: 10.1016/j.media.2007.06.004

Avants BB, Tustison NJ, Song G, Cook PA, Klein A, Gee JC. 2011. A reproducible evaluation of ANTs similarity metric performance in brain image registration. NeuroImage 54:2033–2044. DOI: 10.1016/j.neuroimage.2010.09.025

Awad A, Blomstedt P, Westling G, Eriksson J. 2020. Deep brain stimulation in the caudal zona incerta modulates the sensorimotor cerebello-cerebral circuit in essential tremor. NeuroImage 209:116511. DOI: 10.1016/j.neuroimage.2019.116511

Axer M, Amunts K, Grässel D, Palm C, Dammers J, Axer H, Pietrzyk U, Zilles K. 2011. A novel approach to the human connectome: Ultra-high resolution mapping of fiber tracts in the brain. NeuroImage 54:1091–1101. DOI: 10.1016/j.neuroimage.2010.08.075

Badre D, D’Esposito M. 2009. Is the rostro-caudal axis of the frontal lobe hierarchical? Nature Reviews Neuroscience 10:659–669. DOI: 10.1038/nrn2667

Barthó P, Freund TF, Acsády L. 2002. Selective GABAergic innervation of thalamic nuclei from zona incerta. The European Journal of Neuroscience 16:999–1014. DOI: 10.1046/j.1460-9568.2002.02157.x, PMID: 12383229

Basser PJ, Pierpaoli C. 1996. Microstructural and physiological features of tissues elucidated by quantitative-diffusion-tensor MRI. Journal of Magnetic Resonance. Series B 111:209–219. DOI: 10.1006/jmrb.1996.0086, PMID: 8661285

Behrens TEJ, Berg HJ, Jbabdi S, Rushworth MFS, Woolrich MW. 2007. Probabilistic diffusion tractography with multiple fibre orientations: What can we gain? NeuroImage 34:144–155. DOI: 10.1016/j.neuroimage.2006.09.018

Beliveau V, Ganz M, Feng L, Ozenne B, Højgaard L, Fisher PM, Svarer C, Greve DN, Knudsen GM. 2017. A High-Resolution In Vivo Atlas of the Human Brain’s Serotonin System. The Journal of Neuroscience 37:120–128. DOI: 10.1523/JNEUROSCI.2830-16.2016, PMID: 28053035

Bingham CS, McIntyre CC. 2024. Coupled Activation of the Hyperdirect and Cerebellothalamic Pathways with Zona Incerta Deep Brain Stimulation. Movement Disorders n/a. DOI: 10.1002/mds.29717

Blomstedt P, Persson RS, Hariz G-M, Linder J, Fredricks A, Häggström B, Philipsson J, Forsgren L, Hariz M. 2018. Deep brain stimulation in the caudal zona incerta versus best medical treatment in patients with Parkinson’s disease: a randomised blinded evaluation. Journal of Neurology, Neurosurgery & Psychiatry 89:710–716. DOI: 10.1136/jnnp-2017-317219, PMID: 29386253

Blomstedt P, Sandvik U, Tisch S. 2010. Deep brain stimulation in the posterior subthalamic area in the treatment of essential tremor. Movement Disorders: Official Journal of the Movement Disorder Society 25:1350–1356. DOI: 10.1002/mds.22758, PMID: 20544817

Chometton S, Barbier M, Risold P-Y. 2021. Chapter 11 - The zona incerta system: Involvement in attention and movement. In: Swaab DF, Kreier F, Lucassen PJ, Salehi A, Buijs RM (Eds). *Handbook of Clinical Neurology*, The Human Hypothalamus. Elsevier. p. 173–184. DOI: 10.1016/B978-0-12-820107-7.00011-2

Chometton S, Charrière K, Bayer L, Houdayer C, Franchi G, Poncet F, Fellmann D, Risold PY. 2017. The rostromedial zona incerta is involved in attentional processes while adjacent LHA responds to arousal: c-Fos and anatomical evidence. Brain Structure & Function 222:2507–2525. DOI: 10.1007/s00429-016-1353-3, PMID: 28185007

Chou X, Wang X, Zhang Z, Shen L, Zingg B, Huang J, Zhong W, Mesik L, Zhang LI, Tao HW. 2018. Inhibitory gain modulation of defense behaviors by zona incerta. Nature Communications 9:1151. DOI: 10.1038/s41467-018-03581-6

Coifman RR, Lafon S, Lee AB, Maggioni M, Nadler B, Warner F, Zucker SW. 2005. Geometric diffusions as a tool for harmonic analysis and structure definition of data: Diffusion maps. Proceedings of the National Academy of Sciences 102:7426–7431. DOI: 10.1073/pnas.0500334102

Coudé D, Parent A, Parent M. 2018. Single-axon tracing of the corticosubthalamic hyperdirect pathway in primates. Brain Structure and Function 223:3959–3973. DOI: 10.1007/s00429-018-1726-x

Donahue CJ, Sotiropoulos SN, Jbabdi S, Hernandez-Fernandez M, Behrens TE, Dyrby TB, Coalson T, Kennedy H, Knoblauch K, Essen DCV, Glasser MF. 2016. Using Diffusion Tractography to Predict Cortical Connection Strength and Distance: A Quantitative Comparison with Tracers in the Monkey. Journal of Neuroscience 36:6758–6770. DOI: 10.1523/JNEUROSCI.0493-16.2016, PMID: 27335406

Eggenschwiler F, Kober T, Magill AW, Gruetter R, Marques JP. 2012. SA2RAGE: a new sequence for fast B1+-mapping. Magnetic Resonance in Medicine 67:1609–1619. DOI: 10.1002/mrm.23145, PMID: 22135168

Fischl B. 2012. FreeSurfer. NeuroImage 62:774–781. DOI: 10.1016/j.neuroimage.2012.01.021, PMID: 22248573

FitzGibbon T, Solomon SG, Goodchild AK. 2000. Distribution of calbindin, parvalbumin, and calretinin immunoreactivity in the reticular thalamic nucleus of the marmoset: Evidence for a medial leaflet of incertal neurons. Experimental Neurology 164:371–383. DOI: 10.1006/exnr.2000.7436

Fonov V, Evans A, McKinstry R, Almli C, Collins D. 2009. Unbiased nonlinear average age-appropriate brain templates from birth to adulthood. NeuroImage 47:S102. DOI: 10.1016/S1053-8119(09)70884-5

Fratzl A, Hofer SB. 2022. The caudal prethalamus: Inhibitory switchboard for behavioral control? Neuron 110:2728–2742. DOI: 10.1016/j.neuron.2022.07.018

Garau C, Hayes J, Chiacchierini G, McCutcheon JE, Apergis-Schoute J. 2023. Involvement of A13 dopaminergic neurons in prehensile movements but not reward in the rat. Current Biology 33:4786–4797.e4. DOI: 10.1016/j.cub.2023.09.044

Glasser MF, Coalson TS, Robinson EC, Hacker CD, Harwell J, Yacoub E, Ugurbil K, Andersson J, Beckmann CF, Jenkinson M, Smith SM, Van Essen DC. 2016. A multi-modal parcellation of human cerebral cortex. Nature 536:171–178. DOI: 10.1038/nature18933

Glasser MF, Sotiropoulos SN, Wilson JA, Coalson TS, Fischl B, Andersson JL, Xu J, Jbabdi S, Webster M, Polimeni JR, Van Essen DC, Jenkinson M, WU-Minn HCP Consortium. 2013. The minimal preprocessing pipelines for the Human Connectome Project. NeuroImage 80:105–124. DOI: 10.1016/j.neuroimage.2013.04.127, PMID: 23668970

Guell X, Schmahmann JD, Gabrieli JD, Ghosh SS, Geddes MR. 2020. Asymmetric Functional Gradients in the Human Subcortex. DOI: 10.1101/2020.09.04.283820

Haast RAM, Ivanov D, Uludağ K. 2018. The impact of B1+ correction on MP2RAGE cortical T1 and apparent cortical thickness at 7T. Human Brain Mapping 39:2412–2425. DOI: 10.1002/hbm.24011, PMID: 29457319

Haber SN, Lehman J, Maffei C, Yendiki A. 2023. The Rostral Zona Incerta: A Subcortical Integrative Hub and Potential Deep Brain Stimulation Target for Obsessive-Compulsive Disorder. Biological Psychiatry 93:1010–1022. DOI: 10.1016/j.biopsych.2023.01.006

Hardman C, Ashwell K. 2012. Stereotaxic and Chemoarchitectural Atlas of the Brain of the Common Marmoset (Callithrix jacchus). DOI: 10.1201/b11635

Hollunder B, Ostrem JL, Sahin IA, Rajamani N, Oxenford S, Butenko K, Neudorfer C, Reinhardt P, Zvarova P, Polosan M, Akram H, Vissani M, Zhang C, Sun B, Navratil P, Reich MM, Volkmann J, Yeh F-C, Baldermann JC, Dembek TA, Visser-Vandewalle V, Alho EJL, Franceschini PR, Nanda P, Finke C, Kühn AA, Dougherty DD, Richardson RM, Bergman H, DeLong MR, Mazzoni A, Romito LM, Tyagi H, Zrinzo L, Joyce EM, Chabardes S, Starr PA, Li N, Horn A. 2024. Mapping dysfunctional circuits in the frontal cortex using deep brain stimulation. Nature Neuroscience 27:573–586. DOI: 10.1038/s41593-024-01570-1

Hormigo S, Zhou J, Chabbert D, Sajid S, Busel N, Castro-Alamancos M. 2023. Zona incerta distributes a broad movement signal that modulates behavior. eLife 12:RP89366. DOI: 10.7554/eLife.89366, PMID: 38048270

Horn A, Kühn AA. 2015. Lead-DBS: A toolbox for deep brain stimulation electrode localizations and visualizations. NeuroImage 107:127–135. DOI: 10.1016/j.neuroimage.2014.12.002

Hu T-T, Wang R-R, Du Y, Guo F, Wu Y-X, Wang Y, Wang S, Li X-Y, Zhang S-H, Chen Z. 2019. Activation of the Intrinsic Pain Inhibitory Circuit from the Midcingulate Cg2 to Zona Incerta Alleviates Neuropathic Pain. Journal of Neuroscience 39:9130–9144. DOI: 10.1523/JNEUROSCI.1683-19.2019, PMID: 31604834

Jeurissen B, Descoteaux M, Mori S, Leemans A. 2019. Diffusion MRI fiber tractography of the brain. NMR in Biomedicine 32:e3785. DOI: 10.1002/nbm.3785

Kai J, Khan AR, Haast RA, Lau JC. 2022. Mapping the subcortical connectome using in vivo diffusion MRI: Feasibility and reliability. NeuroImage 262:119553. DOI: 10.1016/j.neuroimage.2022.119553, PMID: 35961469

Kasa LW, Peters T, Mirsattari SM, Jurkiewicz MT, Khan AR, A M Haast R. 2022. The role of the temporal pole in temporal lobe epilepsy: A diffusion kurtosis imaging study. NeuroImage. Clinical 36:103201. DOI: 10.1016/j.nicl.2022.103201, PMID: 36126518

Katsumi Y, Zhang J, Chen D, Kamona N, Bunce JG, Hutchinson JB, Yarossi M, Tunik E, Dickerson BC, Quigley KS, Barrett LF. 2023. Correspondence of functional connectivity gradients across human isocortex, cerebellum, and hippocampus. Communications Biology 6:1–13. DOI: 10.1038/s42003-023-04796-0

Kawana E, Watanabe K. 1981. A cytoarchitectonic study of zona incerta in the rat. Journal Fur Hirnforschung 22:535–541. PMID: 7328309

Kim LH, Lognon A, Sharma S, Tran MA, Chomiak T, Tam S, McPherson C, Eaton SEA, Kiss ZHT, Whelan PJ. 2023. Restoration of locomotor function following stimulation of the A13 region in Parkinson’s mouse models. eLife 12. DOI: 10.7554/eLife.90832

Koechlin E, Summerfield C. 2007. An information theoretical approach to prefrontal executive function. Trends in Cognitive Sciences 11:229–235. DOI: 10.1016/j.tics.2007.04.005

Kreilkamp BAK, Lisanti L, Glenn GR, Wieshmann UC, Das K, Marson AG, Keller SS. 2019. Comparison of manual and automated fiber quantification tractography in patients with temporal lobe epilepsy. NeuroImage: Clinical 24:102024. DOI: 10.1016/j.nicl.2019.102024

Kuhn HW. 1955. The Hungarian method for the assignment problem. Naval Research Logistics Quarterly 2:83–97. DOI: 10.1002/nav.3800020109

Lambert C, Zrinzo L, Nagy Z, Lutti A, Hariz M, Foltynie T, Draganski B, Ashburner J, Frackowiak R. 2012. Confirmation of functional zones within the human subthalamic nucleus: Patterns of connectivity and sub-parcellation using diffusion weighted imaging. NeuroImage 60:83–94. DOI: 10.1016/j.neuroimage.2011.11.082

Larivière S, Bayrak Ş, Vos de Wael R, Benkarim O, Herholz P, Rodriguez-Cruces R, Paquola C, Hong S-J, Misic B, Evans AC, Valk SL, Bernhardt BC. 2023. BrainStat: A toolbox for brain-wide statistics and multimodal feature associations. NeuroImage 266:119807. DOI: 10.1016/j.neuroimage.2022.119807

Lau JC, Xiao Y, Haast RAM, Gilmore G, Uludağ K, MacDougall KW, Menon RS, Parrent AG, Peters TM, Khan AR. 2020. Direct visualization and characterization of the human zona incerta and surrounding structures. Human Brain Mapping 41:4500–4517. DOI: 10.1002/hbm.25137, PMID: 32677751

Levy R. 2024. The prefrontal cortex: from monkey to man. Brain 147:794–815. DOI: 10.1093/brain/awad389

Li J, Bai Y, Liang Y, Zhang Y, Zhao Q, Ge J, Li D, Zhu Y, Cai G, Tao H, Wu S, Huang J. 2022. Parvalbumin Neurons in Zona Incerta Regulate Itch in Mice. Frontiers in Molecular Neuroscience 15:843754. DOI: 10.3389/fnmol.2022.843754, PMID: 35299695

Li J, Peng S, Zhang Y, Ge J, Gao S, Zhu Y, Bai Y, Wu S, Huang J. 2023. Glutamatergic Neurons in the Zona Incerta Modulate Pain and Itch Behaviors in Mice. Molecular Neurobiology 60:5866–5877. DOI: 10.1007/s12035-023-03431-7

Li LX, Li YL, Wu JT, Song JZ, Li XM. 2021. Glutamatergic Neurons in the Caudal Zona Incerta Regulate Parkinsonian Motor Symptoms in Mice. Neuroscience Bulletin 38:1–15. DOI: 10.1007/s12264-021-00775-9, PMID: 34633650

Li Z, Rizzi G, Tan KR. 2021. Zona incerta subpopulations differentially encode and modulate anxiety. Science Advances 7:eabf6709. DOI: 10.1126/sciadv.abf6709, PMID: 34516764

Lin S, Zhu M-Y, Tang M-Y, Wang M, Yu X-D, Zhu Y, Xie S-Z, Yang D, Chen J, Li X-M. 2023. Somatostatin-Positive Neurons in the Rostral Zona Incerta Modulate Innate Fear-Induced Defensive Response in Mice. Neuroscience Bulletin 39:245–260. DOI: 10.1007/s12264-022-00958-y

Liu K, Kim J, Kim DW, Zhang YS, Bao H, Denaxa M, Lim S-A, Kim E, Liu C, Wickersham IR, Pachnis V, Hattar S, Song J, Brown SP, Blackshaw S. 2017. Lhx6-positive GABA-releasing neurons of the zona incerta promote sleep. Nature 548:582–587. DOI: 10.1038/nature23663

Liu M, Blanco-Centurion C, Konadhode R, Begum S, Pelluru D, Gerashchenko D, Sakurai T, Yanagisawa M, van den Pol AN, Shiromani PJ. 2011. Orexin Gene Transfer into Zona Incerta Neurons Suppresses Muscle Paralysis in Narcoleptic Mice. Journal of Neuroscience 31:6028–6040. DOI: 10.1523/JNEUROSCI.6069-10.2011, PMID: 21508228

Lu CW, Harper DE, Askari A, Willsey MS, Vu PP, Schrepf AD, Harte SE, Patil PG. 2021. Stimulation of zona incerta selectively modulates pain in humans. Scientific Reports 11:8924. DOI: 10.1038/s41598-021-87873-w, PMID: 33903611

Maffei C, Girard G, Schilling KG, Aydogan DB, Adluru N, Zhylka A, Wu Y, Mancini M, Hamamci A, Sarica A, Teillac A, Baete SH, Karimi D, Yeh F-C, Yildiz ME, Gholipour A, Bihan-Poudec Y, Hiba B, Quattrone Andrea, Quattrone Aldo, Boshkovski T, Stikov N, Yap P-T, de Luca A, Pluim J, Leemans A, Prabhakaran V, Bendlin BB, Alexander AL, Landman BA, Canales-Rodríguez EJ, Barakovic M, Rafael-Patino J, Yu T, Rensonnet G, Schiavi S, Daducci A, Pizzolato M, Fischi-Gomez E, Thiran J-P, Dai G, Grisot G, Lazovski N, Puch S, Ramos M, Rodrigues P, Prčkovska V, Jones R, Lehman J, Haber SN, Yendiki A. 2022. Insights from the IronTract challenge: Optimal methods for mapping brain pathways from multi-shell diffusion MRI. NeuroImage 257:119327. DOI: 10.1016/j.neuroimage.2022.119327

Maier-Hein KH, Neher PF, Houde J-C, Côté M-A, Garyfallidis E, Zhong J, Chamberland M, Yeh F-C, Lin Y-C, Ji Q, Reddick WE, Glass JO, Chen DQ, Feng Y, Gao C, Wu Y, Ma J, He R, Li Q, Westin C-F, Deslauriers-Gauthier S, González JOO, Paquette M, St-Jean S, Girard G, Rheault F, Sidhu J, Tax CMW, Guo F, Mesri HY, Dávid S, Froeling M, Heemskerk AM, Leemans A, Boré A, Pinsard B, Bedetti C, Desrosiers M, Brambati S, Doyon J, Sarica A, Vasta R, Cerasa A, Quattrone A, Yeatman J, Khan AR, Hodges W, Alexander S, Romascano D, Barakovic M, Auría A, Esteban O, Lemkaddem A, Thiran J-P, Cetingul HE, Odry BL, Mailhe B, Nadar MS, Pizzagalli F, Prasad G, Villalon-Reina JE, Galvis J, Thompson PM, Requejo FDS, Laguna PL, Lacerda LM, Barrett R, Dell’Acqua F, Catani M, Petit L, Caruyer E, Daducci A, Dyrby TB, Holland-Letz T, Hilgetag CC, Stieltjes B, Descoteaux M. 2017. The challenge of mapping the human connectome based on diffusion tractography. Nature Communications 8:1349. DOI: 10.1038/s41467-017-01285-x

Marcus DS, Harwell J, Olsen T, Hodge M, Glasser MF, Prior F, Jenkinson M, Laumann T, Curtiss SW, Van Essen DC. 2011. Informatics and data mining tools and strategies for the human connectome project. Frontiers in Neuroinformatics 5:4. DOI: 10.3389/fninf.2011.00004, PMID: 21743807

Margulies DS, Ghosh SS, Goulas A, Falkiewicz M, Huntenburg JM, Langs G, Bezgin G, Eickhoff SB, Castellanos FX, Petrides M, Jefferies E, Smallwood J. 2016. Situating the default-mode network along a principal gradient of macroscale cortical organization. Proceedings of the National Academy of Sciences 113:12574–12579. DOI: 10.1073/pnas.1608282113, PMID: 27791099

Markello RD, Hansen JY, Liu Z-Q, Bazinet V, Shafiei G, Suárez LE, Blostein N, Seidlitz J, Baillet S, Satterthwaite TD, Chakravarty MM, Raznahan A, Misic B. 2022. neuromaps: structural and functional interpretation of brain maps. Nature Methods 19:1472–1479. DOI: 10.1038/s41592-022-01625-w

Marques JP, Kober T, Krueger G, van der Zwaag W, Van de Moortele P-F, Gruetter R. 2010. MP2RAGE, a self bias-field corrected sequence for improved segmentation and T1-mapping at high field. NeuroImage 49:1271–1281. DOI: 10.1016/j.neuroimage.2009.10.002, PMID: 19819338

May PJ, Basso MA. 2018. Connections Between the Zona Incerta and Superior Colliculus in the Monkey and Squirrel. Brain structure & function 223:371. DOI: 10.1007/s00429-017-1503-2, PMID: 28852862

Mitrofanis J. 2005. Some certainty for the “zone of uncertainty”? Exploring the function of the zona incerta. Neuroscience 130:1–15. DOI: 10.1016/j.neuroscience.2004.08.017, PMID: 15561420

Mitrofanis J, Ashkan K, Wallace BA, Benabid A-L. 2004. Chemoarchitectonic heterogeneities in the primate zona incerta: clinical and functional implications. Journal of Neurocytology 33:429–440. DOI: 10.1023/B:NEUR.0000046573.28081.dd, PMID: 15520528

Mitrofanis J, Mikuletic L. 1999. Organisation of the cortical projection to the zona incerta of the thalamus. The Journal of Comparative Neurology 412:173–185. PMID: 10440718

Monosov IE, Ogasawara T, Haber SN, Heimel JA, Ahmadlou M. 2022. The zona incerta in control of novelty seeking and investigation across species. Current Opinion in Neurobiology 77:102650. DOI: 10.1016/j.conb.2022.102650

Moon HC, Lee YJ, Cho CB, Park YS. 2016. Suppressed GABAergic signaling in the zona incerta causes neuropathic pain in a thoracic hemisection spinal cord injury rat model. Neuroscience Letters 632:55–61. DOI: 10.1016/j.neulet.2016.08.035

Morel A, Magnin M, Jeanmonod D. 1997. Multiarchitectonic and stereotactic atlas of the human thalamus. The Journal of Comparative Neurology 387:588–630. DOI: 10.1002/(sici)1096-9861(19971103)387:4%253C588::aid-cne8%253E3.0.co;2-z, PMID: 9373015

Moriya S, Kuwaki T. 2020. A13 dopamine cell group in the zona incerta is a key neuronal nucleus in nociceptive processing. Neural Regeneration Research 16:1415–1416. DOI: 10.4103/1673-5374.300991, PMID: 33318433

Moriya S, Yamashita A, Masukawa D, Setoyama H, Hwang Y, Yamanaka A, Kuwaki T. 2020. Involvement of A13 dopaminergic neurons located in the zona incerta in nociceptive processing: a fiber photometry study. Molecular Brain 13:60. DOI: 10.1186/s13041-020-00600-w

Mostofi A, Evans JM, Partington-Smith L, Yu K, Chen C, Silverdale MA. 2019. Outcomes from deep brain stimulation targeting subthalamic nucleus and caudal zona incerta for Parkinson’s disease. NPJ Parkinson’s disease 5:17. DOI: 10.1038/s41531-019-0089-1, PMID: 31453317

Mugler III JP, Brookeman JR. 1990. Three-dimensional magnetization-prepared rapid gradient-echo imaging (3D MP RAGE). Magnetic Resonance in Medicine 15:152–157. DOI: 10.1002/mrm.1910150117

Nicolelis MAL, Chapin JK, Lin RCS. 1995. Development of direct GABAergic projections from the zona incerta to the somatosensory cortex of the rat. Neuroscience 65:609–631. DOI: 10.1016/0306-4522(94)00493-O

Ongür D, An X, Price JL. 1998. Prefrontal cortical projections to the hypothalamus in macaque monkeys. The Journal of Comparative Neurology 401:480–505. PMID: 9826274

Oostenveld R, Fries P, Maris E, Schoffelen J-M. 2011. FieldTrip: Open source software for advanced analysis of MEG, EEG, and invasive electrophysiological data. Computational Intelligence and Neuroscience 2011:156869. DOI: 10.1155/2011/156869, PMID: 21253357

Ossowska K. 2020. Zona incerta as a therapeutic target in Parkinson’s disease. Journal of Neurology 267:591–606. DOI: 10.1007/s00415-019-09486-8, PMID: 31375987

Paxinos G, Watson C. 2006. The Rat Brain in Stereotaxic Coordinates: Hard Cover Edition. Elsevier.

Pedregosa F, Varoquaux G, Gramfort A, Michel V, Thirion B, Grisel O, Blondel M, Prettenhofer P, Weiss R, Dubourg V, Vanderplas J, Passos A, Cournapeau D, Brucher M, Perrot M, Duchesnay E. 2011. Scikit-learn: Machine Learning in Python. Journal of Machine Learning Research 12:2825–2830.

Petersen SE, Seitzman BA, Nelson SM, Wig GS, Gordon EM. 2024. Principles of cortical areas and their implications for neuroimaging. Neuron. DOI: 10.1016/j.neuron.2024.05.008

Plaha P, Ben-Shlomo Y, Patel NK, Gill SS. 2006. Stimulation of the caudal zona incerta is superior to stimulation of the subthalamic nucleus in improving contralateral parkinsonism. Brain 129:1732–1747. DOI: 10.1093/brain/awl127

Plaha P, Javed S, Agombar D, O’ Farrell G, Khan S, Whone A, Gill S. 2011. Bilateral caudal zona incerta nucleus stimulation for essential tremor: outcome and quality of life. Journal of Neurology, Neurosurgery, and Psychiatry 82:899–904. DOI: 10.1136/jnnp.2010.222992, PMID: 21285454

Poldrack RA, Kittur A, Kalar D, Miller E, Seppa C, Gil Y, Parker DS, Sabb FW, Bilder RM. 2011. The Cognitive Atlas: Toward a Knowledge Foundation for Cognitive Neuroscience. Frontiers in Neuroinformatics 5. DOI: 10.3389/fninf.2011.00017

Romanowski CA, Mitchell IJ, Crossman AR. 1985. The organisation of the efferent projections of the zona incerta. Journal of Anatomy 143:75–95. PMID: 3870735

Royer J, Paquola C, Valk SL, Kirschner M, Hong S-J, Park B, Bethlehem RAI, Leech R, Yeo BTT, Jefferies E, Smallwood J, Margulies D, Bernhardt BC. 2024. Gradients of Brain Organization: Smooth Sailing from Methods Development to User Community. Neuroinformatics. DOI: 10.1007/s12021-024-09660-y

Saluja S, Qiu L, Wang AR, Campos G, Seilheimer R, McNab JA, Haber SN, Barbosa DAN, Halpern CH. 2024. Diffusion Magnetic Resonance Imaging Tractography Guides Investigation of the Zona Incerta: A Novel Target for Deep Brain Stimulation. Biological Psychiatry. DOI: 10.1016/j.biopsych.2024.02.1004

Schaltenbrand G. 1977. Atlas for stereotaxy of the human brain. Georg Thieme.

Shafiei G, Baillet S, Misic B. 2022. Human electromagnetic and haemodynamic networks systematically converge in unimodal cortex and diverge in transmodal cortex. PLOS Biology 20:e3001735. DOI: 10.1371/journal.pbio.3001735

Shapson-Coe A, Januszewski M, Berger DR, Pope A, Wu Y, Blakely T, Schalek RL, Li P, Wang S, Maitin-Shepard J, Karlupia N, Dorkenwald S, Sjostedt E, Leavitt L, Lee D, Bailey L, Fitzmaurice A, Kar R, Field B, Wu H, Wagner-Carena J, Aley D, Lau J, Lin Z, Wei D, Pfister H, Peleg A, Jain V, Lichtman JW. 2021. A connectomic study of a petascale fragment of human cerebral cortex. bioRxiv 2021.05.29.446289. DOI: 10.1101/2021.05.29.446289

Sharma S, Kim LH, Mayr KA, Elliott DA, Whelan PJ. 2018. Parallel descending dopaminergic connectivity of A13 cells to the brainstem locomotor centers. Scientific Reports 8:7972. DOI: 10.1038/s41598-018-25908-5, PMID: 29789702

Shepherd H, Heartshorne R, Osman-Farah J, Macerollo A. 2024. Dual target deep brain stimulation for complex essential and dystonic tremor - A 5-year follow up. Journal of the Neurological Sciences 457:122887. DOI: 10.1016/j.jns.2024.122887, PMID: 38295533

Sun J, Deng X, Zhu L, Lin J, Chen G, Tang Y, Lu S, Lu Z, Meng Z, Li Y, Zhu Y. 2024. Zona incerta mediates early life isoflurane-induced fear memory deficits. Scientific Reports 14:15136. DOI: 10.1038/s41598-024-66106-w

Sydnor VJ, Larsen B, Bassett DS, Alexander-Bloch A, Fair DA, Liston C, Mackey AP, Milham MP, Pines A, Roalf DR, Seidlitz J, Xu T, Raznahan A, Satterthwaite TD. 2021. Neurodevelopment of the association cortices: Patterns, mechanisms, and implications for psychopathology. Neuron 109:2820–2846. DOI: 10.1016/j.neuron.2021.06.016, PMID: 34270921

Taha A, Gilmore G, Abbass M, Kai J, Kuehn T, Demarco J, Gupta G, Zajner C, Cao D, Chevalier R, Ahmed A, Hadi A, Karat BG, Stanley OW, Park PJ, Ferko KM, Hemachandra D, Vassallo R, Jach M, Thurairajah A, Wong S, Tenorio MC, Ogunsanya F, Khan AR, Lau JC. 2023. Magnetic resonance imaging datasets with anatomical fiducials for quality control and registration. Scientific Data 10:449. DOI: 10.1038/s41597-023-02330-9

Thomas C, Ye FQ, Irfanoglu MO, Modi P, Saleem KS, Leopold DA, Pierpaoli C. 2014. Anatomical accuracy of brain connections derived from diffusion MRI tractography is inherently limited. Proceedings of the National Academy of Sciences 111:16574–16579. DOI: 10.1073/pnas.1405672111

Van Essen DC, Ugurbil K, Auerbach E, Barch D, Behrens TEJ, Bucholz R, Chang A, Chen L, Corbetta M, Curtiss SW, Della Penna S, Feinberg D, Glasser MF, Harel N, Heath AC, Larson-Prior L, Marcus D, Michalareas G, Moeller S, Oostenveld R, Petersen SE, Prior F, Schlaggar BL, Smith SM, Snyder AZ, Xu J, Yacoub E, WU-Minn HCP Consortium. 2012. The Human Connectome Project: a data acquisition perspective. NeuroImage 62:2222–2231. DOI: 10.1016/j.neuroimage.2012.02.018, PMID: 22366334

Venkataraman A, Hunter SC, Dhinojwala M, Ghebrezadik D, Guo J, Inoue K, Young LJ, Dias BG. 2021. Incerto-thalamic modulation of fear via GABA and dopamine. Neuropsychopharmacology 46:1658–1668. DOI: 10.1038/s41386-021-01006-5

Vos de Wael R, Benkarim O, Paquola C, Lariviere S, Royer J, Tavakol S, Xu T, Hong S-J, Langs G, Valk S, Misic B, Milham M, Margulies D, Smallwood J, Bernhardt BC. 2020. BrainSpace: a toolbox for the analysis of macroscale gradients in neuroimaging and connectomics datasets. Communications Biology 3:1–10. DOI: 10.1038/s42003-020-0794-7

Vos de Wael R, Larivière S, Caldairou B, Hong S-J, Margulies DS, Jefferies E, Bernasconi A, Smallwood J, Bernasconi N, Bernhardt BC. 2018. Anatomical and microstructural determinants of hippocampal subfield functional connectome embedding. Proceedings of the National Academy of Sciences 115:10154–10159. DOI: 10.1073/pnas.1803667115

Vu AT, Auerbach E, Lenglet C, Moeller S, Sotiropoulos SN, Jbabdi S, Andersson J, Yacoub E, Ugurbil K. 2015. High resolution whole brain diffusion imaging at 7T for the Human Connectome Project. NeuroImage 122:318–331. DOI: 10.1016/j.neuroimage.2015.08.004, PMID: 26260428

Wang X-Y, Zhang H-Q, Tong K, Han J, Zhao X-Y, Song Y-T, Hao J-R, Sun N, Gao C. 2024. Glutamatergic Projection from the Ventral Tegmental Area to the Zona Incerta Regulates Fear Response. Neuroscience 541:14–22. DOI: 10.1016/j.neuroscience.2024.01.020

Watson C, Lind CRP, Thomas MG. 2014. The anatomy of the caudal zona incerta in rodents and primates. Journal of Anatomy 224:95–107. DOI: 10.1111/joa.12132, PMID: 24138151

Wolters CH, Anwander A, Tricoche X, Weinstein D, Koch MA, MacLeod RS. 2006. Influence of tissue conductivity anisotropy on EEG/MEG field and return current computation in a realistic head model: A simulation and visualization study using high-resolution finite element modeling. NeuroImage 30:813–826. DOI: 10.1016/j.neuroimage.2005.10.014

Wu F-L, Chen S-H, Li J-N, Zhao L-J, Wu X-M, Hong J, Zhu K-H, Sun H-X, Shi S-J, Mao E, Zang W-D, Cao J, Kou Z-Z, Li Y-Q. 2023. Projections from the Rostral Zona Incerta to the Thalamic Paraventricular Nucleus Mediate Nociceptive Neurotransmission in Mice. Metabolites 13:226. DOI: 10.3390/metabo13020226

Yang S, Meng Y, Li J, Li B, Fan Y-S, Chen H, Liao W. 2020. The thalamic functional gradient and its relationship to structural basis and cognitive relevance. NeuroImage 218:116960. DOI: 10.1016/j.neuroimage.2020.116960

Yang Y, Jiang T, Jia X, Yuan J, Li X, Gong H. 2022. Whole-Brain Connectome of GABAergic Neurons in the Mouse Zona Incerta. Neuroscience Bulletin 38:1315–1329. DOI: 10.1007/s12264-022-00930-w, PMID: 35984621

Yarkoni T, Poldrack RA, Nichols TE, Van Essen DC, Wager TD. 2011. Large-scale automated synthesis of human functional neuroimaging data. Nature Methods 8:665–670. DOI: 10.1038/nmeth.1635, PMID: 21706013

Zhang F, Daducci A, He Y, Schiavi S, Seguin C, Smith RE, Yeh C-H, Zhao T, O’Donnell LJ. 2022. Quantitative mapping of the brain’s structural connectivity using diffusion MRI tractography: A review. NeuroImage 249:118870. DOI: 10.1016/j.neuroimage.2021.118870

Zhang L, Zhang P, Qi G, Cai H, Li T, Li M, Cui C, Lei J, Ren K, Yang J, Ming J, Tian B. 2022. A zona incerta-basomedial amygdala circuit modulates aversive expectation in emotional stress-induced aversive learning deficits. Frontiers in Cellular Neuroscience 16.

Zhang X, van den Pol AN. 2017. Rapid binge-like eating and body weight gain driven by zona incerta GABA neuron activation. Science (New York, N.Y.) 356:853–859. DOI: 10.1126/science.aam7100, PMID: 28546212

Zhou H, Xiang W, Huang M. 2021. Inactivation of Zona Incerta Blocks Social Conditioned Place Aversion and Modulates Post-traumatic Stress Disorder-Like Behaviors in Mice. Frontiers in Behavioral Neuroscience 15.

Zhou M, Liu Z, Melin MD, Ng YH, Xu W, Südhof TC. 2018. A Central Amygdala to Zona Incerta Projection is Required for Acquisition and Remote Recall of Conditioned Fear Memory. Nature neuroscience 21:1515–1519. DOI: 10.1038/s41593-018-0248-4, PMID: 30349111

